# Millennial-scale change on a Caribbean reef system that experiences hypoxia

**DOI:** 10.1101/2021.04.06.438665

**Authors:** Blanca Figuerola, Ethan L. Grossman, Noelle Lucey, Nicole D. Leonard, Aaron O’Dea

## Abstract

Coastal hypoxia has become an increasingly acknowledged threat to coral reefs that is potentially intensifying because of increased input of anthropogenic nutrients. Almirante Bay (Caribbean Panama) is a semi-enclosed system that experiences hypoxia in deeper waters which occasionally expand into shallow coral reefs, suffocating most aerobic benthic life. To explore the long-term history of reefs in the bay we extracted reef matrix cores from two reefs that today experience contrasting patterns of oxygenation. We constructed a 1800-year-long record of gastropod assemblages and isotope compositions from six U-Th chronologically-constrained reef matrix cores. We extracted two cores from each reef at 3 m water depth and two additional cores from a deeper part (4.8 m) of the hypoxia-exposed reef. Results show that the deeper part of the hypoxic reef slowed in growth and stopped accreting approximately 1500 years BP while the shallow part of the reef continued to accrete to the present day, in agreement with a model of expanding hypoxia at this time. Our proxy-based approach suggests that differences among these palaeoindicators in the two reefs may have been driven by an increase in hypoxia via eutrophication caused by either natural changes or human impacts. Similar patterns of increasing herbivores and decreasing carbon isotope values occurred in the shallow part of the hypoxic reef during the last few decades. This suggests that hypoxia may be expanding to depths as shallow as 3 m and that shallow reefs are experiencing greater risk due to increased human activity.

## Introduction

Coastal hypoxia, traditionally defined as the depletion of seawater dissolved oxygen (DO) to concentrations less than 2 mg L^-1^, can occur in systems with stratified waters. Oxygen decline is commonly attributed to eutrophication arising from both natural and anthropogenic processes (Diaz and Rosenberg 2008, Rabalais et al. 2010). Excess nutrients can increase algal biomass and fluxes of organic material to bottom waters, fueling microbial respiration that leads to oxygen depletion (Diaz and Rosenberg 2008).

Although increasing hypoxia has long been recognized as a rising global issue for many marine coastal ecosystems (Diaz and Rosenberg 1995, 2008, Breitburg et al. 2018), it was not generally considered as a major threat to coral reefs until relatively recently (Altieri et al. 2017; although see Lapointe and Clark 1992). This perspective may be more a reflection on the relatively poor monitoring of tropical reefs, the apparent ephemerality of hypoxic events on reefs, and the challenges to post-event diagnoses (Altieri et al. 2017, Hughes et al. 2020). Fortuitous monitoring led to the first major hypoxic event being observed in detail on coral reefs in Almirante Bay, Caribbean Panama in 2010 (Altieri et al. 2017). A similar event was also observed in 2017 (Johnson et al. 2018), and detailed monitoring and research on these reefs is documenting the frequency of these events, interpreting the hydrographic processes responsible and unravelling their ecological and evolutionary impacts on reef life.

The 2010 and 2017 hypoxia events documented in Almirante Bay were both episodic and short-lived, lasting less than 5 days. Both resulted in the formation of geographically delineated “dead zones” and widespread damage to the fringing coral reef ecosystems (Fig. 1) in waters deeper than 3–4 m depth (Altieri et al. 2017, Johnson et al. 2018). The Almirante Bay may be naturally prone to hypoxia because it is semi-enclosed with limited inflow from the open ocean (Kaufmann and Thompson 2005), has a low tidal range, is protected from winds, and receives high rainfall. All of these characteristics increase the chance that waters will stratify and surface oxygen will fail to diffuse or become mixed into benthic waters (Diaz and Rosenberg 2008). The increasing eutrophication from runoff of nutrients that has been observed in the region (Seemann et al. 2014), likely caused by deforestation and fertilization from local banana plantations and other human effluents (Stephens 2008, Cramer 2013, Aronson et al. 2014, Graniero et al. 2016, O’Dea et al. 2020), could explain the perceived increased frequency of hypoxia in the bay. Alternatively, episodic hypoxic events may have naturally occurred for millennia, and simply been unobserved due to the lack of long-term proxy data and ephemerality of hypoxia. Long-term environmental and ecological changes have been reconstructed using reef cores (Cramer et al. 2017, 2020) but these were taken on reefs outside the most hypoxic part of the bay (colored red in Fig. 1). Notwithstanding, one of these studies (Cramer et al. 2020) reported an increase in the proportion of infaunal deposit- and suspension-feeding bivalve communities over the last 1000 years and suggested this may be evidence for a gradual decrease in oxygen levels within the reef sediments over time.

**Fig. 1.**
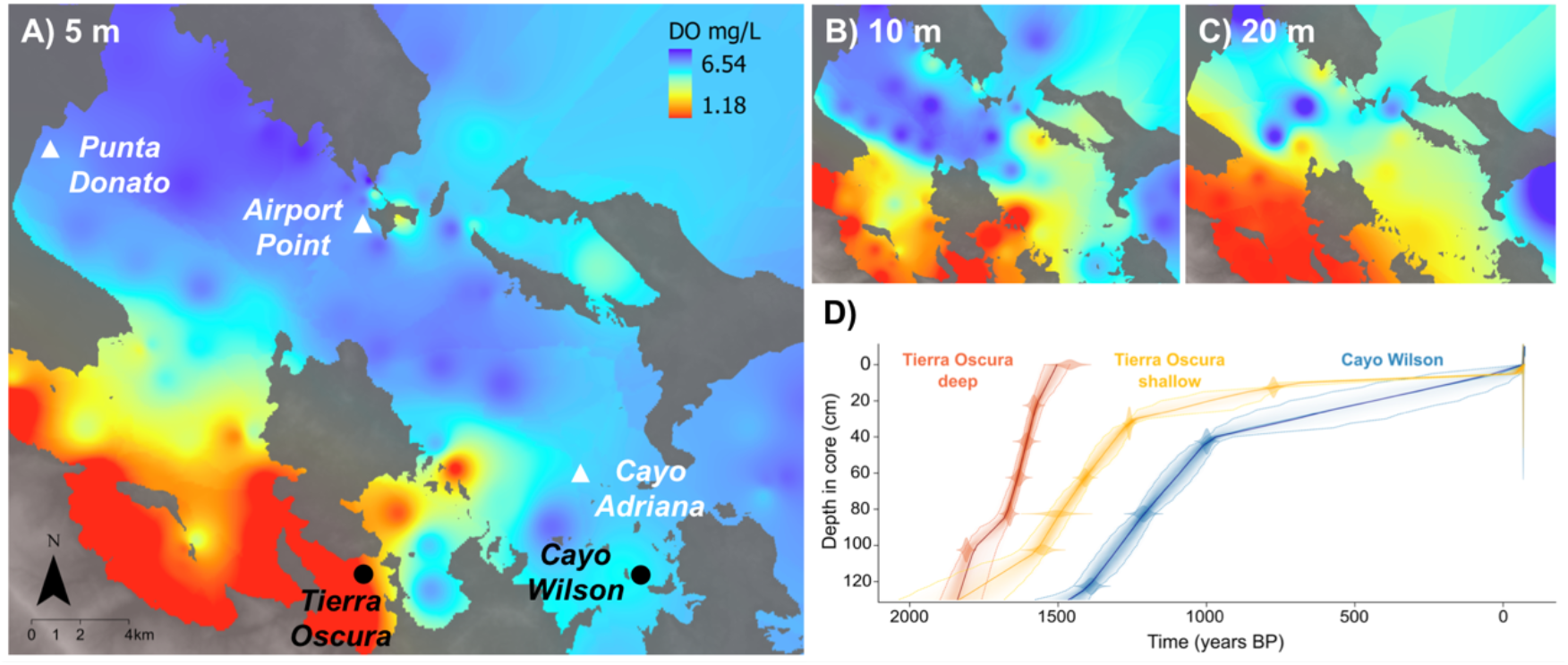
Sampling sites and reef accretion trends. Maps of Almirante Bay showing sites sampled in this and previous studies (black and white, respectively) and the spatial distribution of hypoxia at 5 (A), 10 (B) and 20 m (C) depths (September 2017). DO range at each map: 5 m = 6.54–1.18 mg L^-1^; 10m = 6.24–0.24 mg L^-1^; 20 m: 6.54–0.20 mg L^-1^. (D) Bayesian age-depth models using the R package rbacon (Blaauw and Christen 2011) based on one replicate core at each site and depth. Darker colors indicate more likely calendar ages, dotted lines indicate 95% confidence intervals, and each solid line shows the ‘best fit’ based on the weighted mean age for each depth.

A recent study showed that DO concentration and, to a lesser extent, temperature, are the most important conditions to consider when predicting environmental impacts on a key reef herbivore in the bay (Lucey et al. 2020a). However, there remain uncertainties in the predicted effect of climate change on oxygenation depletion because diverse factors, other than increasing temperature, may interact such as eutrophication, ocean acidification, sea‐level rise and land cover (Altieri and Gedan 2015).

In this study, we quantified the abundances of microgastropods (<5 mm), which are valuable environmental proxies in many marine benthic communities because of their high diversity, and ecological roles, relatively short generation times and good preservation in the fossil record from hundreds to thousands of years (Kidwell and Bosence 1991, Kidwell 2001, Geiger et al. 2007). Microgastropods can also be recovered from cores in substantial numbers, providing high resolution records. Their wide range of feeding modes, life habits, reproductive strategies and life histories leads to a variety of functional roles in reef ecosystems which can provide information on habitats and environments (e.g., Fredston-Hermann et al. 2013) and on the occurrence of taxa without fossil remains (e.g., polychaetes) (Kay et al. 1998).

The oxygen isotopic compositions of mollusc shells (δ^18^O_shell_) are also a powerful archive of past conditions, reflecting the seawater isotopic composition and temperature on daily to millennial timescales (e.g. Schöne et al. 2004, Surge and Barrett 2012, Prendergast and Stevens 2014, Fortunato 2015, Grossman et al. 2019). δ^18^O_shell_ follows a negative quasi-linear relationship with temperature (∼-0.2‰ per °C; Epstein et al. (1953)), but also reflects the δ^18^O of ambient water (δ^18^O_water_), which covaries with salinity (Schmidt 1998). For example, on Panama’s Caribbean coast, δ^18^O_water_ varies from +0.5 to +1.3‰ for a salinity range from 33 to 36 psu (Tao et al. 2013). δ^13^C_shell_ generally reflects the δ^13^C of dissolved inorganic carbon (δ^13^C_DIC_). In shallow environments, δ^13^C_DIC_ varies as a function of photosynthesis, respiration, ocean-atmosphere gas exchange and mixing with ^13^C-depleted freshwater runoff, but can also be influenced by mollusc metabolism (McConnaughey and Gillikin 2008, Strauss et al. 2012).

In this study, we used stable isotopes and gastropod assemblages in reef matrix cores in Almirante Bay to examine environmental and ecological changes over millennia in fringing reefs that experiences hypoxia (Tierra Oscura) and limited hypoxia (Cayo Wilson).

## Methods

### Study area

Almirante Bay is a semi-enclosed coastal lagoon (∼450 km^2^ surface area) bordered by mangroves and delimited by the Bocas del Toro Archipelago (Guzmán et al. 2005). The bay experiences high rainfall (∼3000 mm y^-1^; Paton 2019). Seawater temperatures are typically higher close to the mainland relative to offshore due to the restricted nature of water circulation (Kaufmann and Thompson 2005). Surface salinities of 30 to 34 psu can fall to 20 psu during heavy rain (Kaufmann and Thompson 2005). Tidal range is small (<0.5 m; Fig. 1), and water exchange with the Caribbean Sea occurs through three narrow passages (Kaufmann and Thompson 2005). Due to the local weather conditions and hydrogeomorphology, Almirante Bay experiences seasonal or perennial oxygen depletion in bottom waters, which can extend close to the surface (Lucey et al. 2020b).

The reefs under study, Tierra Oscura and Cayo Wilson, fringe small mangrove-lined cays close to small creeks that experience periodic flooding (Fig. 1). Circulation models suggest flow strengths are comparable, but with higher water retention times in Tierra Oscura than Cayo Wilson (Li and Reidenbach 2014). Tierra Oscura is also located closer to the mainland and therefore likely experiences greater freshwater input and experiences higher water temperatures (Fig. 1). These conditions promote stratification and increased nutrient input, and appear to explain why hypoxic conditions are more common around Tierra Oscura compared to Cayo Wilson (Fig. 1) (Lucey et al. 2020b).

### Environmental measurements

To augment existing environmental data in Almirante Bay (D’Croz et al. 2005, Guzmán et al. 2005, Kaufmann and Thompson 2005, Seemann et al. 2014, Altieri et al. 2017, Lucey et al. 2020b), we made new DO and temperature measurements. We assessed the spatial extent of hypoxia at 5 m, 10 m and approximately 20 m throughout Almirante Bay using a YSI multiparameter sonde (YSI EXO2 & EXO optical DO Smart Sensor) on 26 September 2017 at 83 sites (Fig. 1). Data points were interpolated with Kriging methods using ArcGIS to visualize spatial patterns in DO.

We also deployed optical DO loggers (MiniDOT, PME; YSI Pro2013) at the reef sampling sites and recorded temperature (°C) and DO values every 10 min (precision of ± 0.1°C and ± 5%, respectively) to determine diel variation (2- and 9-month and at Tierra Oscura shallow and deep, respectively, and 3-month deployments at Cayo Wilson) between March and November 2018.

### Sampling and processing

Reef matrix cores were collected from Tierra Oscura and Cayo Wilson in March and June, 2018. We used percussion (push) coring with a 3 m-long aluminium irrigation pipe (7.6 cm external diameter) to extract six cores. At Tierra Oscura, we extracted two replicate cores ∼5 m apart at 2.4 depth through living branching *Porites* reefs and two replicate cores at 4.6 m depth through a dead branching *Porites* matrix (Table S1). At Cayo Wilson, we took two replicate cores at 1.8 m depth on living branching *Porites* reefs. At 10 m depth, there is a sparse reef in Cayo Wilson, while the sediment consists of fine mud and it is covered by anoxic bacterial mats at 10 m depth in Tierra Oscura (authors’ pers. obs.).

The six cores were split lengthwise, with one half archived and the other half sliced into 5 cm sections (hereinafter referred as samples). We analysed every second sample starting from the top (most recent) of each core (e.g. 0–5 cm, 10–15 cm) for a total of 11 or 12 samples per core. Each sample was first dried, weighed, and wet sieved to collect the 2 mm and 500 μm fractions. All gastropod shells in the 2 mm fraction and a split of the 500 μm fraction were picked. Only specimens with a well-preserved apex or aperture (depending on species) were counted. Macro- and microgastropod identifications were based on the available literature (Abbott 1974, Kay et al. 1998, Sasaki 2008, Tunnell 2010) and by comparison with the molluscan reference collection in the Tropical Marine Historical Ecology Laboratory (Smithsonian Tropical Research Institute, Panama).

### Chronology by U-Th dating corals

*Porites* fragments within the cores were U-Th dated (Nu I Plasma Multi-Collector Inductively-Coupled Plasma Mass Spectrometer, MC-ICP-MS; Radiogenic Isotope Facility, University of Queensland), following procedures and analytical protocols previously described (Clark et al. 2012, Cramer et al. 2017) (Supplementary material Appendix 1). From these dates, we estimated accretion rates using linear interpolation between each pair of ages.

### Carbon and oxygen isotope analyses of gastropod shells

Stable isotope analyses were performed on 227 specimens belonging to six common genera (*Anachis, Caecum, Cerithiopsis, Meioceras* and *Modulus*) were selected from the better-dated core at each site (Stable Isotope Geosciences Facility, Texas A&M University; Supplementary material Appendix 1; Fig. S3). We analysed 19 core samples (every fourth sample starting from the top of each core; i.e., e.g., 0–5 cm, 20–25 cm) and analysed at least two replicate specimens from each species. We selected the shells of herbivores that live in near-surface sediments to capture the isotope signal in surface sediments for the non-parametric locally weighted regression (LOESS) and non-metric multidimensional scaling (NMDS) analyses. Mean δ^18^O and δ^13^C values at the three sites and core-top samples were compared using one-way ANOVA. In ANOVA tests, homogeneity of variances was verified with the Levene’s-test. Although the Shapiro-Wilk test revealed that there was a violation of the normality assumption, we still applied ANOVA was still used because it is considered a robust test to violations of normality and a more robust analysis than other non-parametric analyses (e.g., Box 1953, Lix et al. 1996). Whenever a difference was established in the ANOVA tests, multiple comparisons were performed using the Tukey’s HSD test to determine differences between means.

### Temporal trends in gastropod community composition

Relative abundances of gastropod families were calculated for each sample (Supplementary material Appendix 1). Each family was later assigned a life mode (functional group) based on the categories described by Todd (2001) (Table S3). The Pearson correlation was used to test the relationships between isotopes, accretion rates and functional groups. Temporal changes in functional groups and in isotope composition within each core and among the cores were assessed using LOESS-smoothed trend lines using the R package Lattice (Sarkar 2008). Bray-Curtis dissimilarities were calculated and differences in the functional groups between samples and sites were visualized using an NMDS with the metaMDS function (R package Vegan; Oksanen et al. 2019). The data used were log (1 + x) transformed to reduce ordination stress. Two dimensions were calculated (Legendre and Legendre 1998). Next, an analysis of similarity (ANOSIM) was used to test both significant difference between groups of samples and relationships between functional groups and isotope composition using the *envfit* function. The function *envfit* calculates the regression of environmental variables with ordination axes and tests the significance of this regression by permutation test. The matrix of abiotic variables used included carbon and oxygen isotope values. The significance of correlations (*p <* 0.05) between gastropod family and isotope compositions was assessed using 999 random permutations of the abiotic variables. All analyses were performed with the R version 3.5.0 (R Core Team 2019).

## Results

### Chronology and accretion rates of cores

All six cores were composed of a dense framework of branching corals infilled a matrix of sandy-muddy carbonate grains. Visually, branching *Porites* spp. dominated the coral assemblage in all cores, except from the bottom to the 70– 90 cm depth section at Cayo Wilson, where *Agaricia tenuifolia* Dana, 1846 visually dominated at both cores. The twenty ^230^Th ages obtained from the six cores spanned from 1809 to 766 yrs BP (before present, i.e. 1950). ^230^Th age errors (2σ) ranged from ±5 to ±26 yrs (Table S2). The lowest intervals (100 cm) of the two deeper cores from Tierra Oscura yielded ages of 1809 and 1553 yrs BP (cores 1 and 2, respectively), while the sole core-top age (core 1) was 1461 years BP, representing the prehistorical period (Fig. 1D; Table S2). The shallow cores from Tierra Oscura spanned from ∼1500 yrs BP (100 cm) to 814 yrs BP (10 cm). Lastly, Cayo Wilson cores provided an age of 1389 yrs BP at 120 cm and an inferred age of -68 yrs BP at the core-top (inferred age for living coral), representing the prehistorical‐modern (post‐industrial) period. The shift from *Agaricia-* to *Porites*-dominance in the Cayo Wilson site occurred ∼1200 yrs BP. Reef accretion rates at the shallow Tierra Oscura and Cayo Wilson sites peaked at ∼1500 yrs BP and ∼1400 yrs BP, respectively, and slowed earlier in Tierra Oscura than Cayo Wilson (∼1300 yrs BP versus ∼1000 yrs BP) (Fig. 1D).

### Modern environmental data

At Tierra Oscura, temperatures ranged from 27.3°C to 33.5°C at 2.4 m depth and from 29.3°C to 32.7°C at 4.6 m depth over a 9 and 2 month period, respectively. At Cayo Wilson, temperatures ranged from 27.5°C to 31.9°C at 1.8 m over a 3 month period (Fig. S1).

At Tierra Oscura, DO values ranged from 0.02–14.76 mg L^-1^ at 2.4 m depth and 2.18–6.07 mg L^-1^ at 4.6 m depth over a 9 and 2 month period, respectively (Fig. S2). At Cayo Wilson DO ranged from 2.55 to 8.93 mg L^-1^ at 1.8 m depth over a 3 month period (Fig. S2).

DO fluctuated diurnally on both shallow reefs with minimum levels occurring at night (Fig. S2; Table S4). The magnitude of DO fluctuations was highest in shallower waters, and can be attributed to metabolic processes (photosynthesis and respiration) (Lucey et al., 2020). Additionally, the greater variability in DO for Tierra Oscura shallow compared with Cayo Wilson corresponds to severe deep-water hypoxia (Lucey et al. 2020b).

The ranges of temperature and DO generally support the conclusion that hypoxic conditions correspond to warmer temperatures, corroborating previous measures of seasonal conditions in the bay (Lucey et al. 2020b). However, our monitoring data were too short (especially Tierra Oscura deep and Cayo Wilson) and taken during different seasons to detect strong seasonal patterns of hypoxia on the specific reefs and are thus not directly comparable. The spatial analysis showed hypoxic conditions were most severe (1) close to the mainland (Tierra Oscura) compared with the areas closer to the ocean (Cayo Wilson; Fig. 1) and (2) in deeper waters (20 m and 10 m compared with 5 m; Fig. 1A, B, C). These findings are supported by 8 years of monitoring data (Lucey et al. 2020b).

### Isotopic signals and patterns in functional groups

Six sediment cores resulting in 69 samples were analysed for gastropod family composition. Among the 14556 specimens picked, 13269 belonged to 18 families (average of 192.3 specimens/sample; Table S5). Each family was assigned to one of four functional groups (herbivores, carnivores, sponge parasites, parasites on polychaetes and on echinoderms; Table S3). Herbivores dominated all reef environments (68%), followed by parasites (24%) and carnivores (7%; Table S4).

Oxygen and carbon isotopic compositions of micro- and macrogastropods averaged -1.4 ± 0.4‰ and 1.2 ± 0.9‰, respectively. The low variability in δ^18^O reflects the generally narrow range in temperature and salinity. No significant difference in δ^18^O was observed between sites for whole cores (*F* = 0.1513, *df* = 105, *p* = 0.86) or core-tops (*F* = 0.56, *df* = 14, *p* = 0.58). The relatively high variability in δ^13^C reflects the wide range in DO content in response to relative rates of photosynthesis, respiration, and ventilation, superimposed on taxonomic and microhabitat (infaunal) effects. There was no significant difference in mean δ^13^C_shell_ between sites (*F* = 1.72, *df* = 105, *p* = 0.1877); however, the Cayo Wilson core-top has a higher mean δ^13^C_shell_ than core-tops at Tierra Oscura shallow (*p* = 0.002) and Tierra Oscura deep (*p* < 0.001), the trend expected with hypoxia at Tierra Oscura.

At Tierra Oscura deep, δ^13^C_shell_ values were significantly lower in the two most recent samples relative to the older samples (*F* = 12.82, *df* = 31, *p*<0.001; Fig. 2), consistent with decreasing oxygen content. A δ^18^O_shell_ decrease between the recent and older samples was also significant (*F* = 2.855, *df* = 31, *p* = 0.03; Fig. 2). While δ^18^O_shell_ decreased overall up-core in the Tierra Oscura deep core, only the difference between core-top samples and 80 cm section were significant; *p =* 0.02). A significant decrease of δ^13^C_shell_ (*envfit, p =* 0.03) corresponds to a shift to a community dominated by herbivorous taxa from circa 1800 (or ∼1600) to ∼1500 yrs BP (Fig. 3; ANOSIM R = 0.432, *p =* 0.019).

**Fig. 2.**
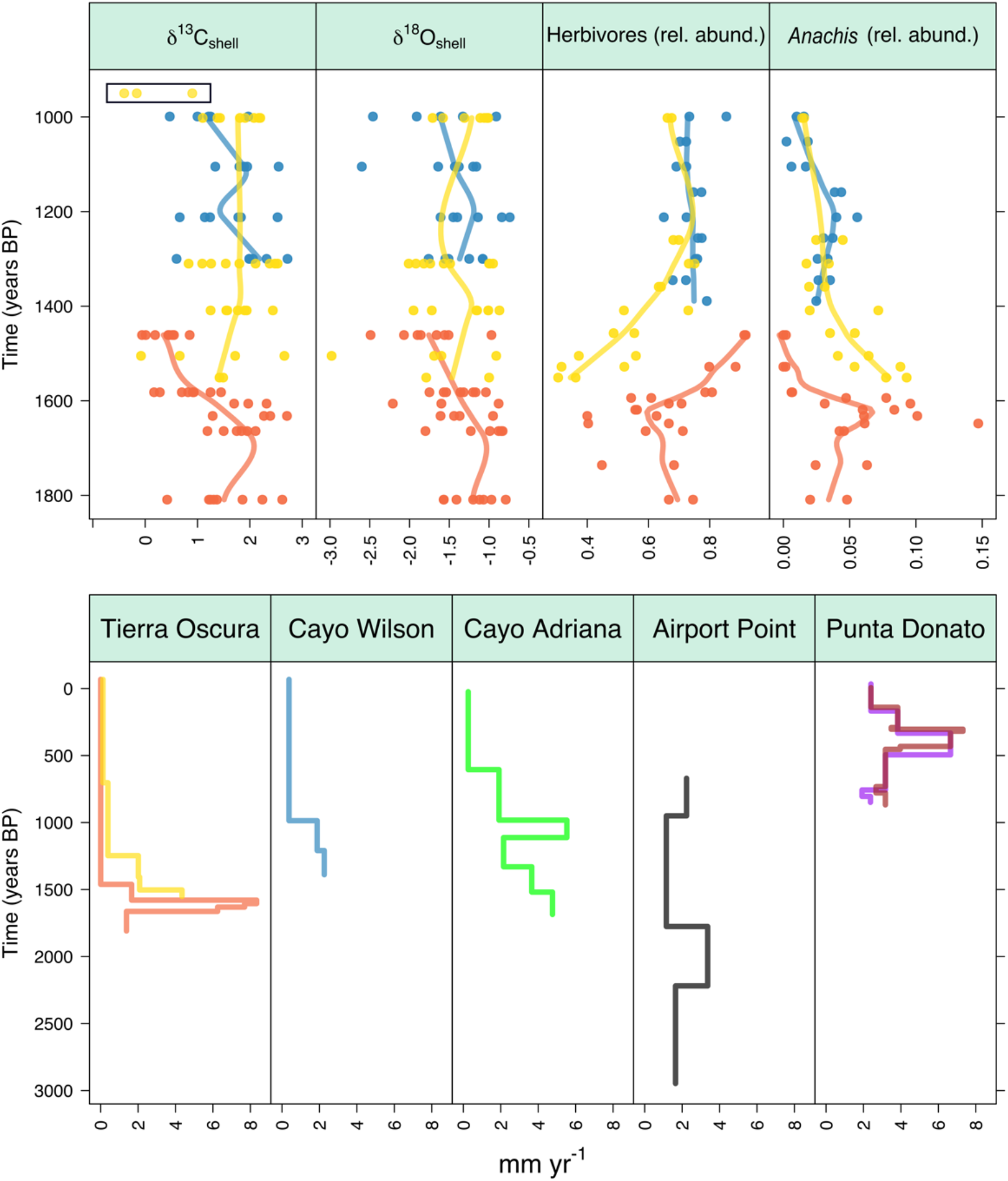
Isotope paleoenvironmental and biotic data in the hypoxic site Tierra Oscura deep (orange), shallow (yellow) and the site Cayo Wilson shallow (blue) from ∼1800 to 1000 yrs BP. Accretion rates in sites sampled in this and previous studies (Cramer et al. 2017, 2020). Recent samples with important trends are indicated by the black rectangle.

**Fig. 3.**
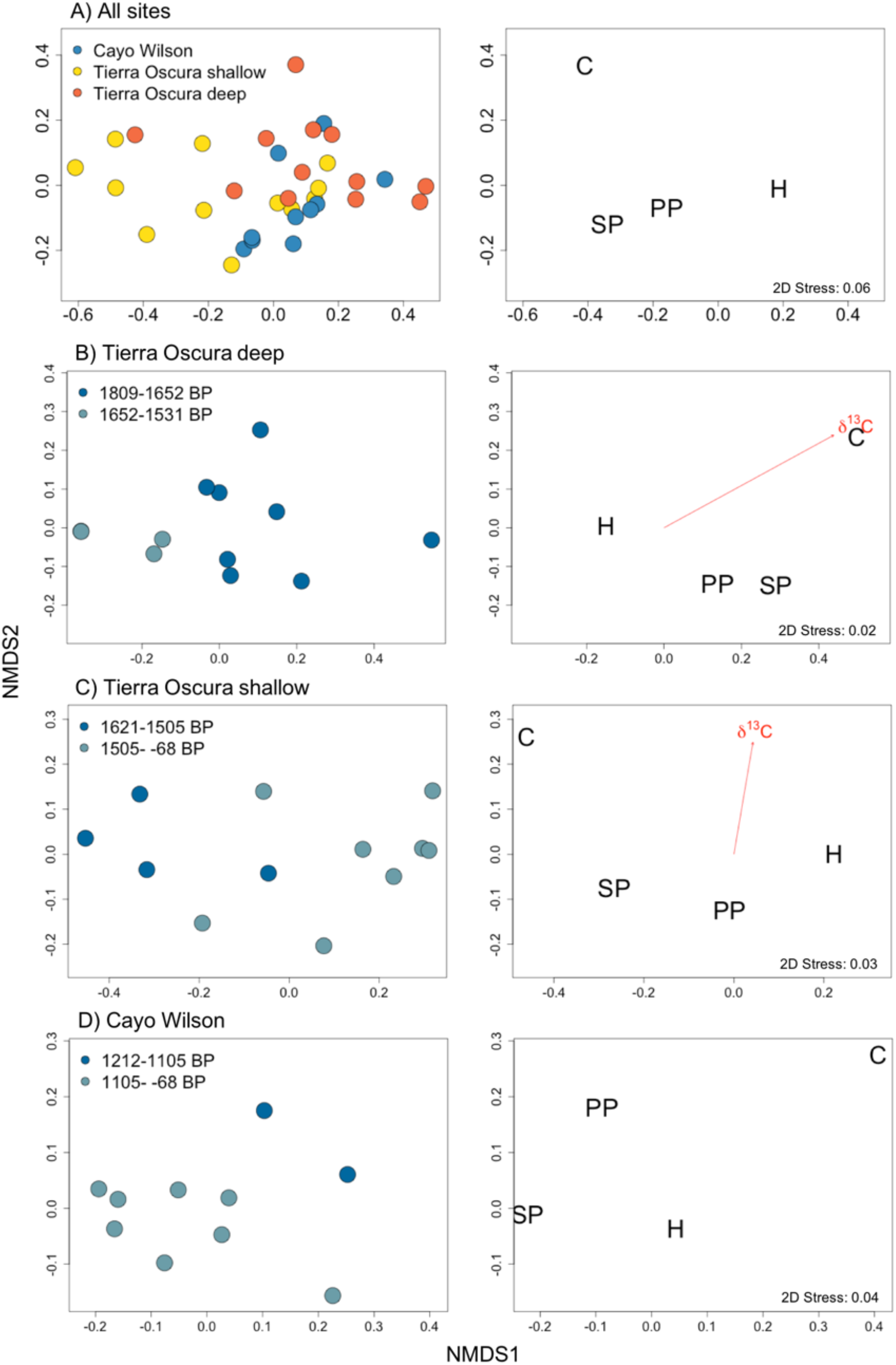
Non-metric multidimensional scaling (NMDS) ordination of functional groups (Stress < 0.10). A) Ordination of samples from all the sites, B) the hypoxic site Tierra Oscura deep, C) shallow and D) the site Cayo Wilson shallow. Significant correlations (*p <* 0.05) were obtained using the *envfit* function for carbon isotope composition and were indicated by vectors (arrow points towards higher values). Samples are color-coded by the sites and years BP. Functional groups: H: herbivores; C: carnivores; SP: sponge parasites; PP: parasites on polychaetes.

At Tierra Oscura shallow, significantly lower δ^13^C_shell_ values were also found in recent relative to older samples (*F* = 3.83, *df* = 26, *p* = 0.009; Fig. 2), but the difference in mean δ^18^O_shell_ values between these samples was not significant (*F* = 1.14, *df* = 26, *p* = 0.36). From ∼1600 (or ∼1500) yrs BP to present, the community shifted from dominance of carnivores to dominance of parasites on polychaetes (Fig. 3; ANOSIM R = 0.579, *p =* 0.009). In addition, relative abundance of herbivores increased from ∼1600 to 1200 yrs ago (Fig. 3). The transition was also characterized by a significant δ^13^C_shell_ decrease (*envfit, p <* 0.017; Fig. 3).

At Cayo Wilson, there was no significant difference in δ^13^C and δ^18^O between samples (δ^13^C_shell_ values: *F* = 0.80, *df* = 48, *p* = 0.55; δ^18^O_shell_: *F* = 0.74, *df* = 32, *p* = 0.60; Fig. 2). From ∼1200–1100 yrs BP to present, the community shifted from carnivore dominance to dominance of herbivores and sponge parasites (Fig. 3; ANOSIM R = 0.6509, *p =* 0.049), coinciding with the brief shift from *Agaricia* to *Porites*. The community change did not correlate with changes in isotope composition (*envfit, p >* 0.05).

The timing of changes in functional groups in the sites rarely coincided with the shift from fast to slow reef accretion rates (Fig. S4). There were strong negative correlations between herbivores and δ^13^C_shell_ (Pearson’s r = -0.83, *p* < 0.001), δ^18^O_shell_ (r = -0.73, *p* < 0.01), sponge parasites (r = -0.82, *p* < 0.001) and carnivores (r = -0.78, *p* < 0.01) in Tierra Oscura deep; and between herbivores and sponge parasites (r = -0.92, *p* < 0.001) and carnivores (r = -0.76, *p* < 0.01) in Tierra Oscura shallow. In Cayo Wilson, there were strong negative correlations between sponge parasites and herbivores (r = -0.79, *p* < 0.01) and accretion rates (r = -0.90, *p* < 0.001) and a strong positive correlation between carnivores and δ^18^O_shell_ (r = 0.87, *p* < 0.01).

## Discussion

We reconstructed environmental and ecological changes over millennia in two Caribbean reefs that today experience contrasting patterns of oxygenation (Fig. 1). In the reef that today experiences hypoxia (Tierra Oscura Deep) we observed increasing herbivore abundances and a decline in isotope values as the deeper portion of the reef transitioned from actively fast accreting to non‐ accreting ∼1500 years BP (Fig. 2).

### Signs of deterioration in coral reef condition

Shallow and deeper reefs experienced different patterns of reef accretion (Fig. 2). The deeper portion of Tierra Oscura was actively accreting at relatively fast rates until ∼1500 yrs BP, when it stopped accreting, suggesting a rapid decline in coral reef health at this time (Figs. 1D and 2). In contrast, the shallow portions of both Tierra Oscura and Cayo Wilson reefs continued to accrete up to today and both had living corals. Accretion rates did decline in the shallow portions but this was likely limited by vertical accommodation space (Spencer 2011) as the reefs approached sea level (Caribbean sea level had more or less stabilized by ∼1500 BP) (Hibbert et al. 2018).

Isotope signals also suggest a prehistoric deterioration of reef health. The deeper portion of Tierra Oscura showed declines in δ^13^C_shell_ values (*p <* 0.05) prior to the cessation of coral growth, supporting evidence of a historical increase in eutrophication, which likely led to the current hypoxic conditions (Fig. 2). Increasing organic input results in enhanced respiration of organic matter through heterotrophic processes in the water column and bottom sediments from coastal environments that produce ^13^C-depleted CO_2_ (i.e., lower δ^13^C values) (i.e., Gooday et al. 2009, Strauss et al. 2012). This depleted δ^13^C_shell_ signal was also observed in the shallow part of the Tierra Oscura reef, although the reef continued to accrete to the present day. The δ^13^C of surface specimens will be influenced by the Suess effect, which can account for a decline of roughly 0.7‰ since 1980 (Gruber et al. 1999). However, this is only half of the shift seen in many taxa (e.g., *Anachis, Meioceras, Modulus*).

Overall, δ^18^O_shell_ trend decreased with time in the Tierra Oscura deep reef, which, in combination with decreasing δ^13^C_shell_, suggests increasing input of freshwater (Fig. 2). However, δ^18^O_shell_ showed no consistent patterns through time in the Tierra Oscura shallow reef. Perhaps a combination of a temporal increase of freshwater input (lower δ^18^O_shell_ values) and cooling of the top layers of the water (higher δ^18^O_shell_ values) buffered the effect of salinity variation on δ^18^O_shell_ in some samples. In particular, Almirante Bay experiences unusual temperature inversions (warmer water at deeper depth) due to a density-driven water column stratification during rainy periods accompanied by, clouds and wind which cool the top layers of water – also registered in this study in September 2018 – which could mask any temporal isotopic signal (Kaufmann and Thompson 2005, Neal et al. 2014).

In this study, all reefs have a past and a current high abundance of low trophic levels (herbivores; including the indicative species *Modulus modulus* (Linnaeus, 1758) of high organic input), which is indicative of high nutrient levels and high algal biomass (Houbrick 1980, Odum 1985, McClanahan 1992, Fichez et al. 2005, Levin et al. 2009, Cramer et al. 2015). However, only the deeper and shallow portion of the Tierra Oscura reef experienced an increase in the relative abundance of herbivores from ∼1600 to 1200 yrs ago, significantly related to decreases of δ^13^C_shell_ (*p <* 0.05). The significantly lower mean δ^18^O_shell_ values of the core-top samples (*p <* 0.05) and the strong negative correlation between oxygen isotope values and herbivores (r = -0.73, *p* < 0.01) in Tierra Oscura deep also suggests a historical increase of freshwater and nutrient levels compared to the shallow part and Cayo Wilson (Fig. 2). The increase of herbivores may be a response to a historical increase of algae (e.g. turf algae), which are more successful with high nutrient input and are becoming increasingly dominant on many Caribbean coral reefs (Bruno et al. 2009, Vermeij et al. 2010).

### Hypoxia as a potential driver of deep reef shutdown

Our findings suggest hypoxia could have been a driver of deep reef shutdown as the declines in δ^13^C_shell_ values occurred earlier in the deeper than the shallow part of the hypoxia-exposed reef. In particular, low oxygen typically occurs close to the seafloor and in deeper portions of stratified water columns, where rates of oxygen consumption are likely to be highest and replenishment lowest (Altieri and Diaz 2019). On the Tierra Oscura reef today, living coral extends down to about 3 m whereupon it shifts sharply into coral rubble which, in places, is covered in mud and bacterial mats. Our historical reconstructions fit the model that this deeper portion of reefs close to the mainland were actively accreting until roughly 1500 years whereupon hypoxia stress led to their shutdown (Fig. 4). By comparing the current spatial distribution of DO in the bay at 5, 10 and 20 m depths (see Fig. 1A, B, C) and available historical data of reef accretion and shutdowns in the bay, we find support for the model of expanding hypoxia from deep to shallow portions over the last millennia. In particular, we found an early decline of the Tierra Oscura deep reef condition (∼1582 yrs BP) prior to the shutdown, compared with the timing of the decline of other more exposed fringing reefs from deeper waters (6–7 m depth) in Almirante Bay (Cayo Adriana and Airport Point, ∼750–650 yrs BP; Punta Donato, ∼450 yrs BP) reported by Cramer et al. (2020) (Figs. 1 and 2).

**Fig. 4.**
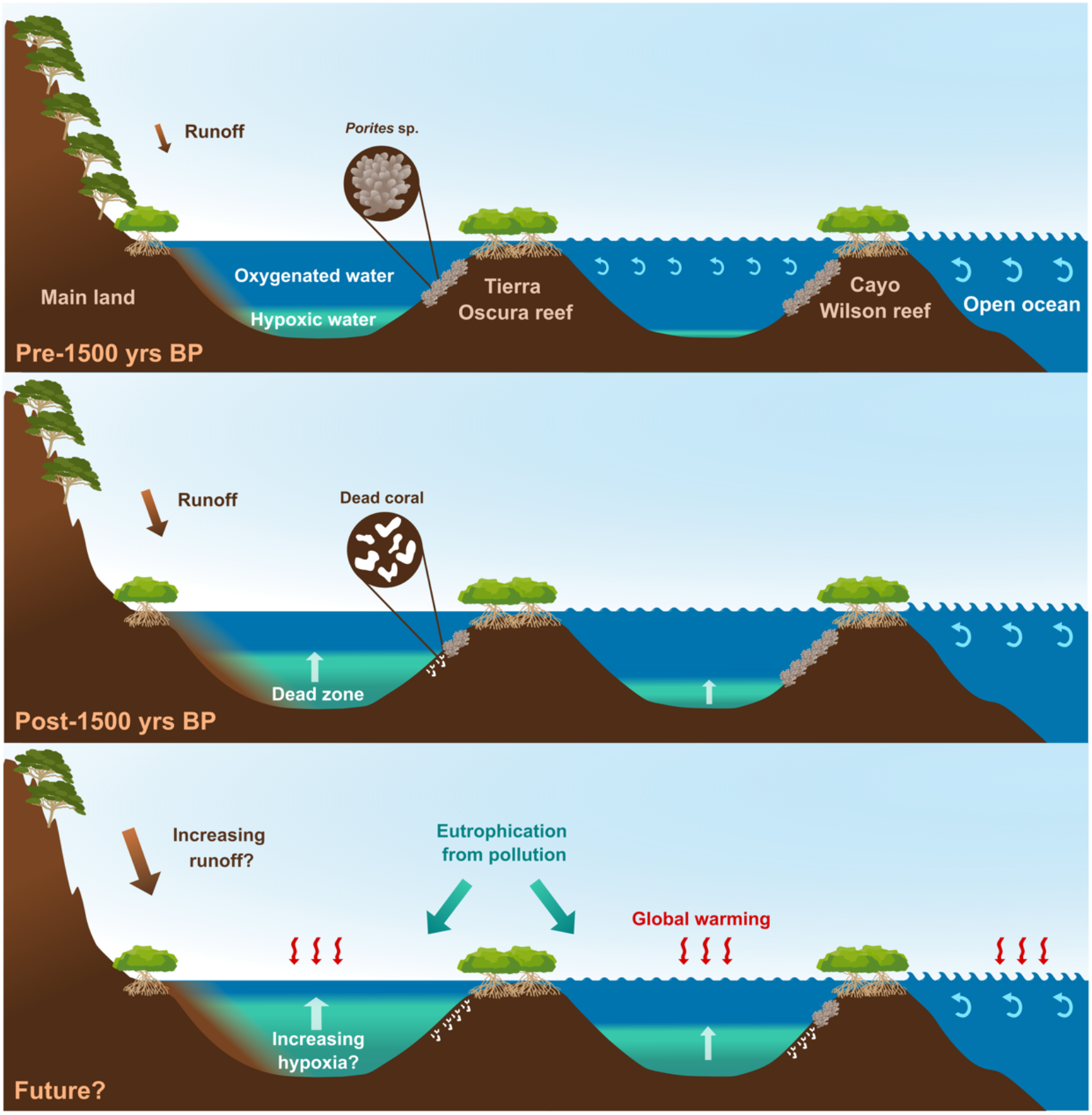
Infographic showing the vertical and horizontal expansion of the hypoxic zone with time in Tierra Oscura and Cayo Wilson. The historical expansion of human populations could result in deforestation on the mainland, initiating an increase in eutrophic and hypoxic conditions, which could lead to the deep reef shutdown in Tierra Oscura approximately 1500 years ago. Shallow reefs are becoming more at risk due to increased anthropogenic activity.

Further, the temporal shift from *Agaricia* to *Porites* at both cores in Cayo Wilson at ∼1200 yrs BP may signal that the conditions could have worsened at this site. These semi-exposed reefs may have drifted toward the hypoxic reef state of more enclosed systems, with a current dominance of *Porites* spp. in shallow waters. *Agaricia* appears to be more affected by high organic carbon and hypoxia than *Porites* (Laboy-Nieves et al. 2001, Kuntz et al. 2005, Altieri et al. 2017). However, more research is needed to confirm this hypothesis as we did not detect any past carbon isotope change.

Some gastropod taxa and functional groups also provide evidence of historical increases of hypoxic conditions. For example, the decline in the relative abundance of *Anachis*, a genus that is particularly sensitive to hypoxic conditions (Díaz Asencio et al. 2016, Kuk-Dzul and Díaz-Castañeda 2016), in the deep and shallow portions of the Tierra Oscura reef from ∼1600–1200 yrs BP corroborates the model of worsening DO concentrations at this reef (Fig. 2). In Tierra Oscura shallow, we also found negative correlations between herbivores and sponge parasites (r = -0.92, *p* < 0.001) and carnivores (r = - 0.76, *p* < 0.01), which suggest similar patterns of change in both portions of the hypoxic reef. For instance, while sponges and polychaetes can grow under stressful conditions and moderate hypoxia (Kay 1980, Pearse 1987, Fichez et al. 2005), these groups are more susceptible to severe hypoxia than other taxa with reported extensive necrosis effects in the case of the sponges (Riedel et al. 2014, Altieri et al. 2017). In addition, the current lack of a nightly decrease in DO in the deeper portion (Fig. S2; Table S4) reflects a simplified ecosystem with low biodiversity and trophic complexity characteristic of a reef that stopped accreting. While our findings support the hypothesis that coastal eutrophication-induced hypoxia is the driver of the reef shutdown, further studies are required to determine other potential causes.

### Natural versus anthropogenic drivers of reef cessation

The earliest known semi-permanent human settlements in the region, radiocarbon dated to ∼3440–2580 yrs BP, are found at Black Creek, 9 km west of Bocas del Toro in Costa Rica (Baldi 2011). However, human populations and their impacts on coastal systems appear to have expanded greatly ∼1350–1250 yrs BP as evidenced from a rapid appearance of shell middens with a wide array of marine resources appearing around Almirante Bay (Linares 1980, Wake and Kay 2013) (Fig. 1). Additionally, by this time conch harvesting was sufficiently intense to have driven declines in size at maturity in *Strombus pugilis* (O’Dea et al. 2014). Large basaltic metates (grinding tools) found on the non-basaltic islands around Almirante Bay, and remains of oceanic fish species from this period (Wake and Kay 2013), demonstrate that humans were able to travel far using large canoes. It is conceivable that the expansion of human populations at this time resulted in deforestation on the mainland, initiating an increase in eutrophic conditions at Tierra Oscura and the shutdown of the deeper portion of the reef there. However, we caution against such a conclusion without further evidence. The human history of the coastal areas remains undefined outside of two small regions in Almirante Bay (Aguacate Peninsula and Sitio Drago), and there is little information available on the human presence on the Caribbean slope of Bocas del Toro mainland at this time. Records from the broader region, however, does support a wider intensification of human impacts on the Isthmus of Panama at this time. In nearby Costa Rica, Taylor (2011) and Taylor et al. (2013) found evidence that agricultural practices peaked from c. 1718 yrs BP to 1400 yrs BP and Anchukaitis and Horn (2005) found evidence of a similar intensification of agriculture c. 1400 yrs BP. If agriculture did significantly expand at this time, the increased erosion and nutrient runoff may have played a role in environmental changes in Almirante Bay. We suggest this hypothesis be explored further using a variety of geochemical and biological proxies for eutrophication such as other stable isotopes (e.g. sedimentary δ^15^N) and preserved remains (e.g. diatom composition), as well as trace elements (Ba, Cd and Zn) and lignin (e.g. Aronson et al. 2014) and lipid biomarkers (see review by Gooday et al. 2009). For instance, Aronson et al. (2014) found peaks in lignin-phenols analyzed from cores extracted in Almirante Bay, which suggest intervals of heavy rainfall.

Non-anthropogenic drivers could also explain the ecosystem changes that occurred in the late Holocene in Almirante Bay. Unfortunately, however, there are few records of natural environmental change from which to test this possibility. High-resolution stalagmite δ^18^O records reveal a significant ENSO-driven shift in average rainfall patterns on the Isthmus of Panama starting around 1500 yrs BP, coincident with the deep reef shutdown. But the shift is from wetter to drier conditions (Lachniet et al. 2004) which would, presumably, have resulted in *less* not *more* run-off from land.

In summary, it is not currently possible to unequivocally assign a driver of the deep-reef shutdown at Tierra Oscura. What limited records exist suggest that shifting rainfall or large-scale ENSO patterns cannot explain the ecosystem changes observed. Deforestation driven by human expansion remains a potential hypothesis that is supported by a mechanism of increased eutrophication driven by land-use changes around Almirante Bay, but a significant dearth of archeological and other evidence means this question currently remains unresolved.

### The future of reefs in Almirante Bay: Warning signs

Our model (Fig. 4) suggests a long term/gradual expansion of the deep-water hypoxic conditions in Almirante Bay towards shallow waters over time. We see evidence of this first in the deeper portion of the Tierra Oscura reef, which is close to the mainland, around c. 1600 yrs BP with changes in the ecological composition of gastropod assemblages and a shift to lower values of δ^13^C_shell_ both indicating increases in organic content and low oxygen levels at this time. These shifts preceded the entire shutdown of the reef, implicating expanding hypoxia as the cause of reef shutdown. The shallow portion of this reef continued to accrete at typical rates, suggesting hypoxic waters did not reach the upper portions of the reef at this time. However, the most recent samples (which we predict span the last 1000 years) reveal a shift toward lower values of δ^13^C_shell_ that parallel the shifts that occurred just prior to the deep reef shutdown. This suggests that low oxygen conditions may be impeaching onto the shallow portions of these reefs to an extent that has not previously occurred. If this is the case, the two intense hypoxic events observed in the bay in the last 10 years (Altieri et al. 2017, Johnson et al. 2018, Lucey et al. 2020b) may indeed be reflecting an intensification of hypoxic conditions with no historical precedence. As agriculture of rainforest within the catchment area and human populations expand, the root causes of eutrophication in the Bay are likely to increase. Our historical data suggest that these reefs can rapidly shift from fast accreting, living reefs to non-accreting “dead” reefs via the vertical expansion of hypoxia, and that these already “marginal” reefs (Schlöder et al. 2013) may be approaching an environmental threshold if hypoxic conditions continue to expand. Controlling the likely drivers of hypoxia, such as land clearance, fertilizer use and human population expansion may help, but the processes leading to increased hypoxia are likely to also be exacerbated by global warming (Breitburg et al. 2018).

## Acknowledgments

We thank Jorge Morales, Ramiro Solís, Brígida de Gracia, Lucía Rodríguez and Jarrod Scott for help field and logistical support and four reviewers for their input and constructive suggestions. We would also like to thank Dr. Chris Maupin, laboratory manager at the Stable Isotope Geosciences Facility at Texas A&M University for helping in the isotope analyses. This project, the postdoctoral fellowship of B. F. and her research visit to Texas A&M University were financially supported by the *Secretaría Nacional de Ciencia y Tecnología e Innovación* (SENACYT; 47-2017-4-FID16-239), *Sistema Nacional de Investigación* (SENACYT), the Scholarly Studies Program (Smithsonian Institution), Smithsonian Tropical Research Institute, Texas A&M University and donations from M. Selin and family, J. Bilyk, V. and B. Anders, J. and M. Bytnar and the Young Presidents Organization (Los Angeles Gold Chapter).

## Notes

### Competing Interest Statement

The authors have declared no competing interest.

## References

Abbott, R. T. 1974. American seashells: the marine mollusks of the Atlantic and Pacific coasts of North America. - New York, NY: Van Nostrand Reinhold Company.

Altieri, A. H. and Gedan, K. B. 2015. Climate change and dead zones. - Glob. Change Biol. 21: 1395–1406.

Altieri, A. H. and Diaz, R. J. 2019. Chapter 24 - Dead Zones: Oxygen Depletion in Coastal Ecosystems. - In: Sheppard, C. (ed), World Seas: an Environmental Evaluation (Second Edition). Academic Press, pp. 453– 473.

Altieri, A. H. et al. 2017. Tropical dead zones and mass mortalities on coral reefs. - Proc. Natl. Acad. Sci. U. S. A. 114: 3660–3665.

Anchukaitis, K. J. and Horn, S. P. 2005. A 2000-year reconstruction of forest disturbance from southern Pacific Costa Rica. - Palaeogeogr. Palaeoclimatol. Palaeoecol. 221: 35–54.

Aronson, R. B. et al. 2014. Land use, water quality, and the history of coral assemblages at Bocas del Toro, Panamá. - Mar. Ecol. Prog. Ser. 504: 159–170.

Baldi, N. F. 2011. Explotación temprana de recursos costeros en el sitio Black Creek (4000-2500 A.P.), Caribe sur de Costa Rica. - Rev. Arqueol. Am.: 85–121.

Blaauw, M. and Christen, J. A. 2011. Flexible paleoclimate age-depth models using an autoregressive gamma process. - Bayesian Anal. 6: 457–474.

Box 1953. Non-normality and tests on variances. - Biometrika 40: 318–335.

Breitburg, D. et al. 2018. Declining oxygen in the global ocean and coastal waters. - Science 359: eaam7240.

Bruno, J. F. et al. 2009. Assessing evidence of phase shifts from coral to macroalgal dominance on coral reefs. - Ecology 90: 1478–1484.

Clark, T. R. et al. 2012. Spatial variability of initial 230Th/232Th in modern Porites from the inshore region of the Great Barrier Reef. - Geochim. Cosmochim. Acta 78: 99–118.

Cramer, K. L. 2013. History of human occupation and environmental change in Western and Central Caribbean Panama. - Bull. Mar. Sci. 89: 955–982.

Cramer, K. L. et al. 2015. Molluscan subfossil assemblages reveal the long- term deterioration of coral reef environments in Caribbean Panama. - Mar. Pollut. Bull. 96: 176–187.

Cramer, K. L. et al. 2017. Prehistorical and historical declines in Caribbean coral reef accretion rates driven by loss of parrotfish. - Nat. Commun. 8: 14160.

Cramer, K. L. et al. 2020. Millennial-scale change in the structure of a Caribbean reef ecosystem and the role of human and natural disturbance. - Ecography 43: 283–293.

D’Croz, L. et al. 2005. The effect of fresh water runoff on the distribution of dissolved inorganic nutrients and plankton in the Bocas Del Toro Archipelago, Caribbean Panama. - Caribb. J. Sci. 41: 414–429.

Diaz, R. J. and Rosenberg, R. 1995. Marine benthic hypoxia: a review of its ecological effects and the behavioural responses of benthic macrofauna. - Oceanogr. Mar. Biol. Annu. Rev. 33: 245–303.

Diaz, R. J. and Rosenberg, R. 2008. Spreading dead zones and consequences for marine ecosystems. - Science 321: 926–929.

Díaz Asencio, L. et al. 2016. Two-year temporal response of benthic macrofauna and sediments to hypoxia in a tropical semi-enclosed bay (Cienfuegos, Cuba). - Rev. Biol. Trop. 64: 177–188.

Epstein, S. et al. 1953. Revised carbonate-water isotopic temperature scale. - Geol. Soc. Am. Bull. 64: 1315–1326.

Fichez, R. et al. 2005. A review of selected indicators of particle, nutrient and metal inputs in coral reef lagoon systems. - Aquat. Living Resour. 18: 125–147.

Fortunato, H. 2015. Mollusks: tools in environmental and climate research*. - Am. Malacol. Bull. 33: 310–324.

Fredston-Hermann, A. L. et al. 2013. Marked ecological shifts in seagrass and reef molluscan communities since the mid-Holocene in the southwestern Caribbean. - Bull. Mar. Sci. 89: 983–1002.

Geiger, D. L. et al. 2007. Techniques for collecting, handling, preparing, storing and examining small molluscan specimens. - Molluscan Res. 27(1): 1–45.

Gooday, A. J. et al. 2009. Historical records of coastal eutrophication-induced hypoxia. - Biogeosciences 6: 1707–1745.

Graniero, L. E. et al. 2016. Stable isotopes in bivalves as indicators of nutrient source in coastal waters in the Bocas del Toro Archipelago, Panama. - PeerJ 4: e2278.

Grossman, E. L. et al. 2019. Freshwater input, upwelling, and the evolution of Caribbean coastal ecosystems during formation of the Isthmus of Panama. - Geology 47: 857–861.

Gruber, N. et al. 1999. Spatiotemporal patterns of carbon-13 in the global surface oceans and the oceanic suess effect. - Glob. Biogeochem. Cycles 13: 307–335.

Guzmán, H. M. et al. 2005. A Site Description of the CARICOMP Mangrove, Seagrass and Coral Reef Sites in Bocas del Toro, Panama. - Caribb. J. Sci. 41: 430–440.

Hibbert, F. D. et al. 2018. A database of biological and geomorphological sea- level markers from the Last Glacial Maximum to present. - Sci. Data 5: 180088.

Houbrick, R. S. 1980. Observations on the anatomy and life history of Modulus modulus (Prosobranchia: Modulidae). - Malacologia 20: 117–142.

Hughes, D. J. et al. 2020. Coral reef survival under accelerating ocean deoxygenation. - Nat. Clim. Change 10: 296–307.

Johnson, M. D. et al. 2018. Shallow-water hypoxia and mass mortality on a Caribbean coral reef. - Bull. Mar. Sci. 94: 143–144.

Kaufmann, K. W. and Thompson, R. C. 2005. Water temperature variation and the meteorological and hydrographic environment of Bocas del Toro, Panama. - Caribb. J. Sci. 41: 392–413.

Kay, E. A. 1980. Micromollusks: techniques and patterns in benthic marine communities. pp. 93–112, in: Water Resources Research Center and Sea Grant, University of Hawaii at Manoa, and Hawaii Water Pollution Control Association. Environmental Survey Techniques for Coastal Water Conference Proceedings. Honolulu, Hawaii.

Kay, E. A. et al. 1998. Mollusca: the southern synthesis. Part B. - In: Fauna of Australia Volume 5. CSIRO Publishing, pp. 565–1234.

Kidwell, S. M. 2001. Preservation of species abundance in marine death assemblages. - Science 294: 1091–1094.

Kidwell, S. M. and Bosence, D. W. J. 1991. Taphonomy and time-averaging of marine shelly faunas, p. 115–209. In Allison, P.A.and Briggs, D.E.G. (eds.) Taphonomy. Releasing the data locked in the fossil record. Plenum, New York.

Kuk-Dzul, J. G. and Díaz-Castañeda, V. 2016. The relationship between mollusks and oxygen concentrations in Todos Santos Bay, Baja California, Mexico. - J. Mar. Biol. 2016: e5757198.

Kuntz, N. M. et al. 2005. Pathologies and mortality rates caused by organic carbon and nutrient stressors in three Caribbean coral species. - Mar. Ecol. Prog. Ser. 294: 173–180.

Laboy-Nieves, E. N. et al. 2001. Mass mortality of tropical marine communities in Morrocoy, Venezuela. - Bull. Mar. Sci. 68: 163–179.

Lachniet, M. S. et al. 2004. A 1500-year El Niño/Southern oscillation and rainfall history for the isthmus of Panama from speleothem calcite. - J. Geophys. Res. Atmospheres in press.

Lapointe, B. E. and Clark, M. W. 1992. Nutrient inputs from the watershed and coastal eutrophication in the Florida keys. - Estuaries 15: 465–476.

Legendre, P. and Legendre, L. 1998. Numerical ecology. - Elsevier, Amsterdam.

Levin, L. A. et al. 2009. Effects of natural and human-induced hypoxia on coastal benthos. - Biogeosciences 6: 2063–2098.

Li, A. and Reidenbach, M. A. 2014. Forecasting decadal changes in sea surface temperatures and coral bleaching within a Caribbean coral reef. - Coral Reefs 33: 847–861.

Linares, O. F. 1980. Ecology and prehistory of the Aguacate Peninsula in Bocas del Toro. - In: Linares, O.F., Ranere, A.J., editors. Adaptive radiations in prehistoric Panama. Peabody Museum Monographs 5. Cambridge, MA: Harvard University Press., pp. 57–66.

Lix, L. M. et al. 1996. Consequences of assumption violations revisited: A quantitative review of alternatives to the one-way analysis of variance F test. - Rev. Educ. Res. 66: 579–619.

Lucey, N. et al. 2020a. Multi-stressor Extremes Found on a Tropical Coral Reef Impair Performance. - Front. Mar. Sci. in press.

Lucey, N. M. et al. 2020b. Oxygen-mediated plasticity confers hypoxia tolerance in a corallivorous polychaete. - Ecol. Evol. 10: 1145–1157.

McClanahan, T. R. 1992. Epibenthic gastropods of the middle Florida keys: the role of habitat and environmental stress on assemblage composition. - J. Exp. Mar. Biol. Ecol. 160: 169–190.

McConnaughey, T. A. and Gillikin, D. P. 2008. Carbon isotopes in mollusk shell carbonates. - Geo-Mar. Lett. 28: 287–299.

Neal, B. P. et al. 2014. When depth is no refuge: cumulative thermal stress increases with depth in Bocas del Toro, Panama. - Coral Reefs 33: 193– 205.

O’Dea, A. et al. 2014. Evidence of size-selective evolution in the fighting conch from prehistoric subsistence harvesting. - Proc. R. Soc. B Biol. Sci. 281: 20140159.

O’Dea, A. et al. 2020. Defining variation in pre-human ecosystems can guide conservation: An example from a Caribbean coral reef. - Sci. Rep. 10: 2922.

Odum, E. P. 1985. Trends expected in stressed ecosystems. - BioScience 35: 419–422.

Oksanen, J. et al. 2019. Vegan: community ecology Ppackage. R package version 2.5-6.

Paton, S. 2019. Bocas del Toro, platform tower_precipitation. The Smithsonian Institution.

Pearse, J. S. 1987. Development, metamorphosis, and seasonal abundance of embryos and larvae of the Antarctic sea urchin.: 126–135.

Prendergast, A. L. and Stevens, R. E. 2014. Molluscs (isotopes): analyses in environmental archaeology. - In: Smith, C. (ed), Encyclopedia of Global Archaeology. Springer, pp. 5010–5019.

Rabalais, N. N. et al. 2010. Dynamics and distribution of natural and human- caused hypoxia. - Biogeosciences 7: 585–619.

Riedel, B. et al. 2014. Effect of hypoxia and anoxia on invertebrate behaviour: ecological perspectives from species to community level. - Biogeosciences 11: 1491–1518.

Sarkar, D. 2008. Lattice: multivariate data visualization with R. Springer, New York. - Use R in press.

Sasaki, T. 2008. Micromolluscs in Japan: taxonomic composition, habitats, and future topics. - Zoosymposia 1: 147–232.

Schlöder, C. et al. 2013. Benthic Community Recovery from Small-Scale Damage on Marginal Caribbean Reefs: An Example from Panama. - Bull. Mar. Sci. 89: 1003–1013.

Schmidt, G. A. 1998. Oxygen-18 variations in a global ocean model. - Geophys. Res. Lett. 25: 1201–1204.

Schöne, B. R. et al. 2004. Sea surface water temperatures over the period 1884–1983 reconstructed from oxygen isotope ratios of a bivalve mollusk shell (Arctica islandica, southern North Sea). - Palaeogeogr. Palaeoclimatol. Palaeoecol. 212: 215–232.

Seemann, J. et al. 2014. Assessing the ecological effects of human impacts on coral reefs in Bocas del Toro, Panama. - Environ. Monit. Assess. 186: 1747–1763.

Spencer, T. 2011. Accommodation Space. - In: Hopley, D. (ed), Encyclopedia of Modern Coral Reefs: Structure, Form and Process. Springer Netherlands, pp. 2–3.

Stephens, C. S. 2008. Outline of history in the province of Bocas del Toro, Panama. Eustis, FL: SFS Publications.

Strauss, J. et al. 2012. 100 Years of benthic foraminiferal history on the inner Texas shelf inferred from fauna and stable isotopes: Preliminary results from two cores. - Cont. Shelf Res. 38: 89–97.

Surge, D. and Barrett, J. H. 2012. Marine climatic seasonality during medieval times (10th to 12th centuries) based on isotopic records in Viking Age shells from Orkney, Scotland. - Palaeogeogr. Palaeoclimatol. Palaeoecol. 350–352: 236–246.

Tao, K. et al. 2013. Quantifying upwelling and freshening in nearshore tropical American environments using stable isotopes in modern gastropods. in press.

Taylor, Z. 2011. Spatial variation in organic carbon and stable isotope composition of lake sediments at Laguna. University of Tennessee, PhD dissertation.

Taylor, Z. et al. 2013. Maize pollen concentrations in Neotropical lake sediments as an indicator of the scale of prehistoric agriculture. - The Holocene 23: 78–84.

Todd, J. A. 2001. Introduction to molluscan life habits database. London, UK: the Natural History Museum. Available from: https://nmita.rsmas.miami.edu/database/mollusc/mollusclifestyles.htm.

Tunnell, J. W. 2010. Encyclopedia of Texas seashells: identification, ecology, distribution, and history. - Texas A&M University Press.

Vermeij, M. J. A. et al. 2010. The effects of nutrient enrichment and herbivore abundance on the ability of turf algae to overgrow coral in the Caribbean. - PLOS ONE 5: e14312.

Wake, T. A. and Kay, M. 2013. Archaeological investigations provide late Holocene baseline ecological data for Bocas del Toro, Panama. - Bull. Mar. Sci. 89: 1015–1035.

