## Supplemental material for "Millennial-scale change on a Caribbean reef system that experiences hypoxia"

This document contains:

Appendix 1

Supplementary Tables S1-S5

Supplementary Figures S1-S5

#### Appendix 1

##### Methods

###### Chronology by U-Th dating corals

### We selected *Porites* spp. fragments, which were identifiable (a proxy for taphonomic preservation). About 150 mg of coral were crushed in an agate mortar and pestle to sand-size chips. and then soaked in 15% H_2_O_2_ overnight. Samples were then ultrasonicated and rinsed in Milli-Q H_2_O (18.2Ω) multiple times to remove organics and surficial detritus. Cleaned samples were then hand-picked under a binocular microscope selecting for only the cleanest aragonite (i.e. no detritus, discoloration or secondary cements). Samples were spiked with a mixed ^229^Th–^233^U tracer dissolved in double-distilled 7N HNO_3_ in ultra-cleaned beakers. To ensure no remnant organics remained in the sample, 6–7 drops of 30% H_2_O_2_ were added to the digested samples which were capped tightly and placed on a hotplate set at 140°C overnight to facilitate complete sample-tracer homogenization. U and Th were separated using conventional anion-exchange column chemistry using Bio-Rad AG 1-X8 resin. Following MC ICP-MS measurement ^230^Th ages were calculated using Isoplot 3.75 Program (Ludwig 2012) using the decay constants of Cheng et al. (2000) and corrected for non-radiogenic ^230^Th contributions using the two-component-mixing model of Clark et al. (2014). At each site, three to five U-Th ages were obtained along one replicate core. From these cores, accretion rates were estimated as achieved previously by dividing the length of an interval by its timespan (c.f. Cramer et al. 2017, 2020). Two ages were obtained from the middle and bottom of the second replicate cores to verify that both cores had the same accumulation rates. The corals from the tops of the shallow cores were not dated as reefs had living coral.

###### Carbon and oxygen isotope analyses of gastropod shells

Prior to analysis, well-preserved gastropod shells were cleaned of mud with deionized water and epiphytic material was removed by carefully scraping the sample. The entire shells or the growing edges of the gastropods, respectively, were homogenized with a mortar and pestle and roughly 40–100 μg of the powdered shells were analyzed for δ^18^O and δ^13^C using a Thermo Scientific MAT 253 isotope ratio mass spectrometer (IRMS) coupled to a Kiel IV automated carbonate reaction system at the Stable Isotope Geosciences Facility at Texas A&M University (<https://stableisotope.tamu.edu/>). Carbon and oxygen isotope analyses were calibrated using the IAEA-603 standard and reported versus VPDB. Isotopic ratios were reported in parts per thousand (per mil, ‰) difference in the isotopic ratio relative to the carbonate standard VPDB. Data are expressed as δ (‰) = (R_x_–R_std_)/R_std_ x 1000 where R_x_ is the isotope ratio of the samples (^18^O/^16^O or ^13^C/^12^C) and R_std_ is the isotope ratio of the standard. Precision was 0.03‰ for δ^13^C and 0.05‰ for δ^18^O.

Six common genera from one replicate core at each site and depth were chosen for isotope analyses. Herbivore shells that can capture the isotope signal in surface sediments (gastropods with a near surface sediment habitat) were selected for the analyses to avoid the microhabitat effect (i.e., incorporation of ^13^C-depleted dissolved inorganic carbon (DIC) from pore waters) (Grossman 1987) which can complicate interpretation of δ^13^C data. For example, *Caecum* shows interspecific variability in its life mode (e.g. Tunnell 2010), *Cerithiopsis* is a parasite, the epiphytic gastropod *Anachis* spp. seem to be very active at night (Kitting 1985).

###### Temporal trends in micro- and macrogastropods

Relative abundances (RA = p*100/n, where *p* is the number of individuals of a family and *n* is the total number of individuals in each sample), were determined for each family. We used relative abundance because changes in sedimentation rates can affect total abundance. In addition, proportional abundances of living molluscan communities are generally accurately represented in death assemblages (Kidwell 2013) as also confirmed by previous studies in Almirante Bay (Cramer et al. 2015, 2020). We computed species proportions from the fraction total weight and fraction total individuals. As the results were similar, we presented the latter.

#### Supplementary Tables

**Supplementary Table S1.** Data information of samples.

| **Site** | **Core** | **Longitude** | **Latitude** | **Water depth (m)** | **Penetration depth (m)** | **Recovery (m)** |
| --- | --- | --- | --- | --- | --- | --- |
| Tierra Oscura deep | AT18-1-1 | -82.25555 | 9.20863 | 4.6 | 2.05 | 1.12 |
| Tierra Oscura deep | AT18-1-2 | -82.25555 | 9.20863 | 4.6 | 2 | 1.32 |
| Tierra Oscura shallow | BF18-3-1 | -82.25675 | 9.20721 | 2.4 | 2.16 | 1.17 |
| Tierra Oscura shallow | BF18-3-2 | -82.25675 | 9.20721 | 2.4 | 2.2 | 1.12 |
| Cayo Wilson | BF18-4-1 | -82.15578 | 9.20474 | 1.85 | 2.32 | 1.14 |
| Cayo Wilson | BF18-4-2 | -82.15578 | 9.20474 | 1.85 | 2.21 | 1.22 |

**Supplementary Table S2.** MC-ICP-MS ^230^Th ages of coral fragments obtained from reef matrix cores collected in the Caribbean. Ratios in parentheses are activity ratios calculated from atomic ratios using decay constants of Cheng et al. (2000). All values have been corrected for laboratory procedural blanks. All errors reported in this table are quoted as 2σ. Uncorrected ^230^Th ages were calculated using Isoplot/EX 3.0 program (Ludwig 2012). ^230^Th (†) ages were corrected using the two-component correction method of Clark et al. (2014) using ^230^Th/^232^Th_hyd_ and ^230^Th/^232^Th_det_ activity ratios of 1.08 ± 0.23 and 0.62 ± 0.14, respectively.

| **Core** | **Core position (mm from top)** | **Sample weight (g)** | **U (ppm)** | **^232^Th (ppb)** | **(^230^Th/ ^232^Th)** | **(^230^Th/^238^U)** | **(^234^U/ ^238^U)** | **corr. ^230^Th Age (ka)** | **Date of chemistry** | **Year (BP)** |
| --- | --- | --- | --- | --- | --- | --- | --- | --- | --- | --- |
| Tierra Oscura deep | 0 | 0.15957 | 2.8781 ± 0.0011 | 1.0045 ± 0.010 | 140.30 ± 051 | 0.016139 ± 0.000056 | 1.1468 ± 0.0011 | 1.531 ± 0.006 | 2020.2 | 1461 ± 6.1 |
|  | 20 | 0.14385 | 2.5778 ± 0.0014 | 2.9391 ± 0.0042 | 46.747 ± 0.1679 | 0.0003758 ± 0.0000018 | 1.1491 ± 0.0012 | 1.6521 ± 0.0080 | 2019.75 | 1582 ± 8.0 |
|  | 40 | 0.15276 | 2.5173 ± 0.0008 | 2.8065 ± 0.0021 | 48.3133 ± 0.2361 | 0.0177519 ± 0.000086 | 1.1458 ± 0.00010 | 1.6753 ± 0.0100 | 2019.2 | 1606 ± 10 |
|  | 60 | 0.1211 | 2.9437 ± 0.0026 | 4.5775 ± 0.0058 | 35.391 ± 0.1367 | 0.018138 ± 0.000068 | 1.1535 ± 0.0020 | 1.7015 ± 0.0097 | 2019.75 | 1632 ± 9.7 |
|  | 80 | 0.14919 | 2.883 ± 0.0009 | 3.1604 ± 0.0031 | 50.8136 ± 0.1402 | 0.018355 ± 0.000048 | 1.1470 ± 0.0006 | 1.7328 ± 0.0070 | 2019.2 | 1664 ± 7 |
|  | 100 | 0.15068 | 2.7218 ± 0.0008 | 1.7454 ± 0.0014 | 93.3848 ± 0.2663 | 0.019736 ± 0.000054 | 1.1447 ± 0.0012 | 1.8784 ± 0.0066 | 2019.2 | 1809 ± 7 |
| Tierra Oscura deep (replicate) | 40 | 0.15068 | 2.783 ± 0.0011 | 3.3421 ± 0.0027 | 44.6021 ± 0.1496 | 0.017653 ± 0.00005 | 1.1447 ± 0.0008 | 1.6662 ± 0.0080 | 2019.2 | 1597 ± 8 |
|  | 100 | 0.15039 | 3.0632 ± 0.0014 | 2.0139 ± 0.0028 | 78.9393 ± 0.2096 | 0.017104 ± 0.000039 | 1.1456 ± 0.0009 | 1.6227 ± 0.0053 | 2019.2 | 1553 ± 5 |
| Tierra Oscura shallow | 10 | 0.15275 | 2.6033 ± 0.0007 | 4.6109 ± 0.0044 | 15.2833 ± 0.0822 | 0.008922 ± 0.000047 | 1.1452 ± 0.0009 | 0.8134 ± 0.0091 | 2019.2 | 814 ± 9 |
|  | 30 | 0.06931 | 3.1801 ± 0.0008 | 19.332 ± 0.015 | 7.55 ± 0.04 | 0.015120 ± 0.000085 | 1.1464 ± 0.0013 | 1.329 ± 0.026 | 2018.5 | 1330 ± 26 |
|  | 60 | 0.15003 | 2.7731 ± 0.001 | 4.0648 ± 0.0027 | 32.6481 ± 0.1115 | 0.015772 ± 0.000047 | 1.1454 ± 0.0008 | 1.4784 ± 0.0084 | 2019.2 | 1409 ± 8 |
|  | 80 | 0.15495 | 2.8321 ± 0.0012 | 5.5471 ± 0.0052 | 26.1295 ± 0.0684 | 0.016868 ± 0.000042 | 1.1461 ± 0.0010 | 1.5738 ± 0.0095 | 2019.2 | 1505 ± 9 |
|  | 100 | 0.15068 | 2.6123 ± 0.0008 | 7.9041 ± 0.0074 | 17.6137 ± 0.0622 | 0.017564 ± 0.000060 | 1.1461 ± 0.0008 | 1.6203 ± 0.0139 | 2019.2 | 1551 ± 14 |
| Tierra Oscura shallow (replicate) | 40 | 0.15154 | 2.8977 ± 0.0010 | 8.7076 ± 0.0092 | 14.477 0.0723 | 0.014338 ± 0.000070 | 1.1460 0.0011 | 1.3103 ± 0.0142 | 2019.2 | 1241 ± 14 |
|  | 100 | 0.14415 | 2.5123 ± 0.0017 | 8.8016 ± 0.0106 | 15.3353 0.0666 | 0.017707 ± 0.000075 | 1.1473 0.0010 | 1.6231 ± 0.0163 | 2019.2 | 1554 ± 16 |
| Cayo Wilson | 40 | 0.14049 | 3.5011 ± 0.0025 | 8.661 ± 0.0119 | 14.3538 0.0583 | 0.011703 ± 0.000045 | 1.1454 0.0012 | 1.0683 ± 0.0112 | 2019.2 | 999 ± 11 |
|  | 80 | 0.15344 | 2.7247 ± 0.0006 | 1.9191 ± 0.0119 | 58.4911 0.2149 | 0.01358 ± 0.000048 | 1.1456 0.0008 | 1.2812 ± 0.0060 | 2019.2 | 1212 ± 6 |
|  | 120 | 0.15248 | 2.8453 ± 0.0009 | 2.0208 ± 0.0016 | 65.9012 0.2243 | 0.015426 ± 0.000051 | 1.1466 0.0010 | 1.4581 ± 0.0063 | 2019.2 | 1389 ± 6 |
| Cayo Wilson (replicate) | 30 | 0.13653 | 2.9972 ± 0.0016 | 4.8873 ± 0.0041 | 16.93 ± 0.07 | 0.009099 ± 0.000040 | 1.1461 ± 0.0007 | 0.8343 ± 0.0081 | 2018.5 | 766 ± 8 |
|  | 100 | 0.13081 | 3.0709 ± 0.0010 | 3.9879 ± 0.003 | 31.670 0.1101 | 0.0135544 ± 0.000046 | 1.1460 0.0008 | 1.2677 ± 0.0074 | 2019.2 | 1199 ± 7 |

**Supplementary Table S3.** Descriptions of the life habitat and feeding categories used to categorize the gastropod families studied by functional groups. Modified from Todd et al. (2001) and Fredston-Hermann et al. (2013).

| **Family** | **Habitat** | **Feeding type** | **Life mode** | **Diet from Todd (2000)** | **Basic diet** | **References** |
| --- | --- | --- | --- | --- | --- | --- |
| Areneidae | under rocks or around hard substrates, shallow waters |  | E | HR | H | http://fossilworks.org; Beesley et al. 1998; Tunnell et al. 2010 |
| Bullidae |  |  | I | HP | H | http://fossilworks.org; Beesley et al. 1998; Tunnell et al. 2010 |
| Caecidae | Caecidae live on a variety of substrata including algae and seagrasses, beneath rocks and rubble and interstitially in gravel, sometimes on intertidal beaches | microphagous feeders on unicellular animals that live on sand grains or small pebbles | E/I | HR/HP | H | http://fossilworks.org; Beesley et al. 1998; Tunnell et al. 2010 |
| Cerithiidae | sandy, coral rubble, on rocks and sand flats | algivores/detritivores | E | HM/HR | H | http://fossilworks.org; Beesley et al. 1998; Tunnell et al. 2010 |
| Cerithiopsidae |  | Ectoparasites feeding usually on sponges on which they live | P | CB | SP | http://fossilworks.org; Beesley et al. 1998; Tunnell et al. 2010 |
| Columbellidae |  | carnivores | E | CP | C | http://fossilworks.org; Beesley et al. 1998; Tunnell et al. 2010 |
| Cylichnidae |  | carnivores feeding on sand-living invertebrates, worms and other mollusks | I | CP | C | http://fossilworks.org; Beesley et al. 1998; Tunnell et al. 2010 |
| Eulimidae |  | ectoparasites feeding primary on echinoderms | P | CB | PE | http://fossilworks.org; Beesley et al. 1998; Tunnell et al. 2010 |
| Fissurellidae |  | herbivores/ carnivores feeding on sponges and other detritivores | E | HR/CB | H | http://fossilworks.org; Beesley et al. 1998; Tunnell et al. 2010 |
| Lottiidae |  | herbivores on microscopic plants growing on rocks | E | HR | H | http://fossilworks.org; Beesley et al. 1998; Tunnell et al. 2010 |
| Marginellidae | rocky reefs to soft shores | carnivores | E | CB | C | http://fossilworks.org; Beesley et al. 1998; Tunnell et al. 2010 |
| Modulidae | shallow, seagrass beds, coral boulders | herbivores feeding on small plants and detritus | E | HR/HP | H | http://fossilworks.org; Beesley et al. 1998; Tunnell et al. 2010 |
| Neritidae |  | detritivores/herbivores feeding on algae | E | HO/HR | H | http://fossilworks.org; Beesley et al. 1998; Tunnell et al. 2010 |
| Pyramidellidae |  | ectoparasites feeding on body fluids of variety of invertebrates, but mainly polychaetes and other molluscs | P | CB | PP | http://fossilworks.org; Beesley et al. 1998; Tunnell et al. 2010 |
| Rissoinidae | sand, under rocks, on algae or marine plants | Biofilm grazer feeding on diatoms and pieces of algae | E | HP | H | http://fossilworks.org; Beesley et al. 1998; Tunnell et al. 2010 |
| Tornidae |  | biofilm grazer feeding on detritus and microscopic plant | E |  | H | http://fossilworks.org; Beesley et al. 1998; Tunnell et al. 2010 |
| Triphoridae | under rocks and dead coral slabs, shallow water or deep | ectoparasites feeding on parasites | E | CB | SP | http://fossilworks.org; Beesley et al. 1998; Tunnell et al. 2010 |
| Turridae | in general >12 m but all depths, most soft substrate | carnivores feeding on polychaetes, sipunculans, nemerteans | E | CP | C | http://fossilworks.org; Beesley et al. 1998; Tunnell et al. 2010 |

**Life mode**

E: epifaunal; I: infaunal; P: parasite.

**Diet**

CP: predatory carnivores. Predators feeding on and killing whole sedentary and mobile macro-organisms and also selective ingesters of foraminifera (foraminiferivores). Included here are scavengers, which with just a few known exceptions, are also predators, shifting facultatively when carrion is present (Britton & Morton 1994).

CB: browsing carnivores. Predators which feed on sedentary, and typically clonal, animals (e.g. corals and other cnidarians, sponges, ascidians) without killing them.This also includes those ‘parasites’, which are ectoparasitic upon mostly relatively larger sedentary or mobile prey. For our purposes, I believe there is no useful distinction between these categories, given the varying host specificities of parasites (e.g. within eulimids, epitoniids and pyramidellids); and a seemingly complete gradation in relative sizes of parasite and host.

HO: herbivorous omnivores. Browsing macroherbivores with unselective omnivory, typically of epifauna attached to macroalgae.

HM: herbivores on fine-grained substrates. Microalgivores, detritivores, microphages and unselective deposit feeder. Also included here is a miscellany of herbivorous non-HR and HP categories, including those living on wood or mangrove substrates.

HR: herbivores on rock, rubble or coral substrates. Microalgivores

HP: herbivores on plant or algal substrates. Micro-and macroalgivores and detritivores on macroalgal and seagrass substrates.

SU: suspension feeders. Includes taxa feeding solely or dominantly upon suspended particles, including mucociliary feeders. Feeding type

**Basic diet**

H: herbivores

C: carnivores

SP: sponge parasites

PP: parasites on polychaetes

PE: parasites on echinoderms

**Supplementary Table S4.** Descriptive statistics of dissolved oxygen (DO) conditions in mg L^-1^.

|  | **Day** |  |  |  |  | **Night** |  |  |  |  |
| --- | --- | --- | --- | --- | --- | --- | --- | --- | --- | --- |
| **Site** | **Mean** | **Min** | **Max** | **Avg max** | **Avg min** | **Mean** | **Min** | **Max** | **Avg max** | **Avg min** |
| Tierra Oscura deep | 3.49 | 2.19 | 5.18 | 3.99 | 3.03 | 3.85 | 2.37 | 6.07 | 4.39 | 3.24 |
| Tierra Oscura shallow | 5.60 | 0.02 | 14.76 | 8.04 | 3.13 | 4.95 | 0.07 | 12.02 | 7.24 | 3.07 |
| Cayo Wilson | 6.25 | 3.32 | 8.94 | 7.27 | 4.76 | 5.71 | 2.55 | 7.30 | 6.46 | 4.39 |

**Supplementary Table S5**. Absolute abundance of functional groups in Tierra Oscura deep, shallow and Cayo Wilson.

| **Functional groups** | **Tierra Oscura deep** | **Tierra Oscura shallow** | **Cayo Wilson** | **Total** | **Percentage** |
| --- | --- | --- | --- | --- | --- |
| Herbivores | 2632 | 2438 | 4012 | 9082 | 68.45 |
| Carnivores | 287 | 355 | 305 | 947 | 7.14 |
| Sponge parasites | 383 | 883 | 1033 | 2299 | 17.33 |
| Parasites on polychaetes | 220 | 349 | 310 | 879 | 6.62 |
| Parasites on echinoderms | 3 | 10 | 49 | 62 | 0.47 |

#### Supplementary Figures

**Supplementary Figure S1.** Time-series of temperature (°C) in Tierra Oscura deep (d) and shallow (s) (2018), and Cayo Wilson (2019).

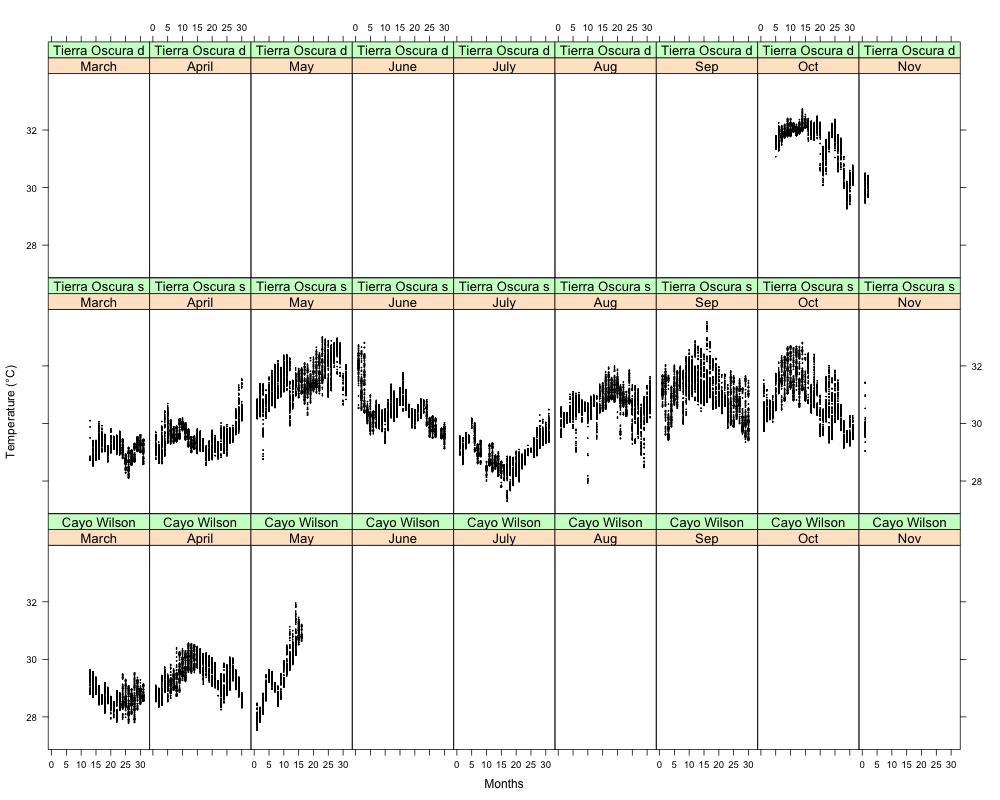

**Supplementary Figure S2.** Time-series of dissolved oxygen (DO) in Tierra Oscura deep (d) and shallow (s) (2018), and Cayo Wilson (2019). The dotted line indicates the threshold of hypoxia.

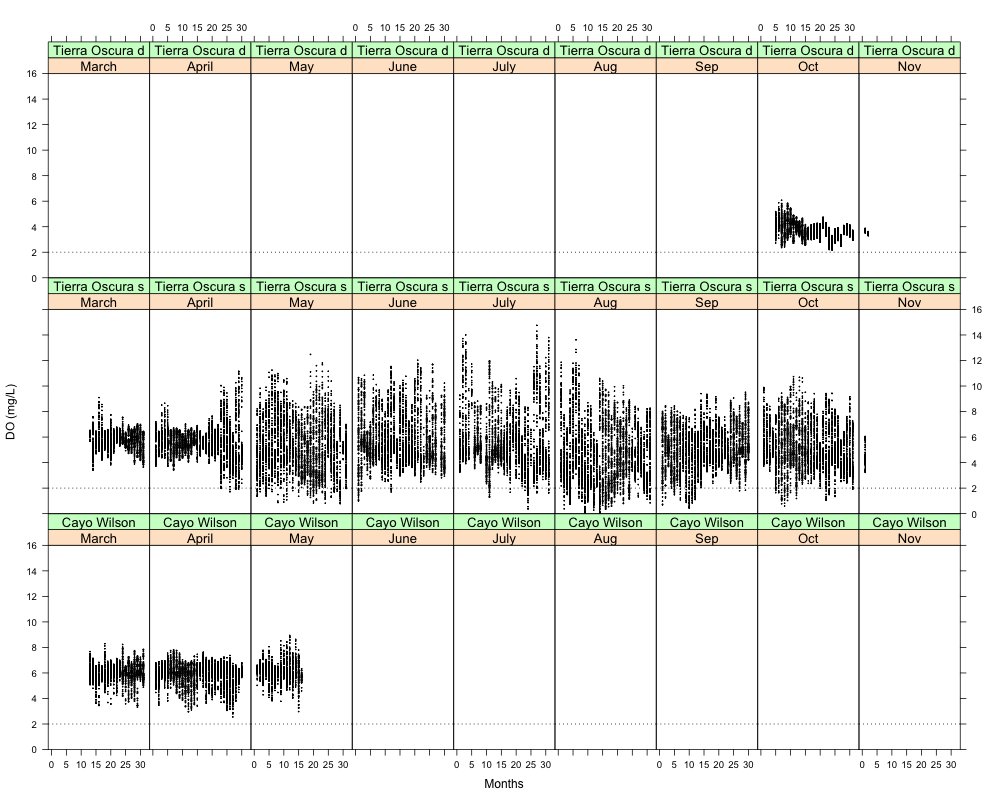

**Supplementary Figure S3.** Temporal trends in δ^13^C_shell_ and δ^18^O_shell_ in common taxa at hypoxic site Tierra Oscura (deep, orange; shallow, yellow) and the site Cayo Wilson site (blue) from a total of 227 specimens (19 samples) selected from one replicate core at each site and depth for isotope analyses. A) Temporal trends in δ^13^C_shell_ of all taxa. The increased spacing between samples in the last 1000 years reflects a dramatic decrease in sedimentation rate. B) Temporal trends in δ^18^O_shell_ of all taxa. Significantly higher δ^13^C_shell_ values were found for the epiphytic gastropod *Anachis* and the parasite *Cerithiopsis* and other taxa (*F* = 23.59, *df* = 231, *p* = 2.2e^-16^; between *Anachis* and *Caecum*, *Cerithiopsis* (p < 0.001) and *Meioceras* (p = 0.002) and between *Cerithiopsis* and *Caecum*, *Meioceras* and *Modulus* (p < 0.001)), supporting evidence of a microhabitat effect (i.e., uptake of ^13^C-depleted DIC from pore waters).

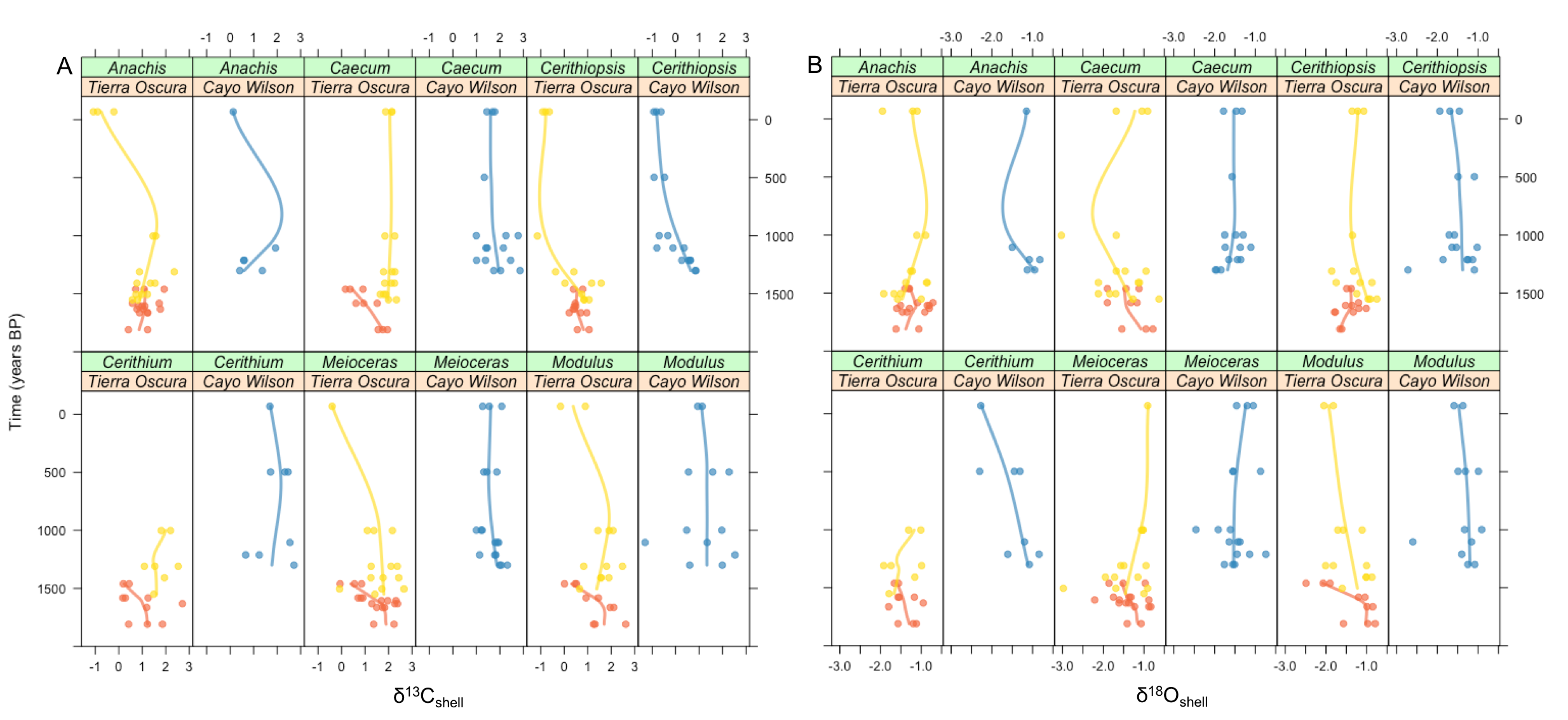

**Supplementary Figure S4.** Correlation matrix of data of Tierra Oscura deep and shallow and Cayo Wilson. The graph provides scatter plots with fitted lines, histograms with kernel density estimations and correlation coefficients (r) with results of significant testing. Accretion: accretion rates; C: carnivores; H: herbivores; PP: parasites on polychaetes; SP: sponge parasites. Significant levels for Pearson correlations: *** *p* < 0.001, ** *p* < 0.01, * *p* < 0.05.

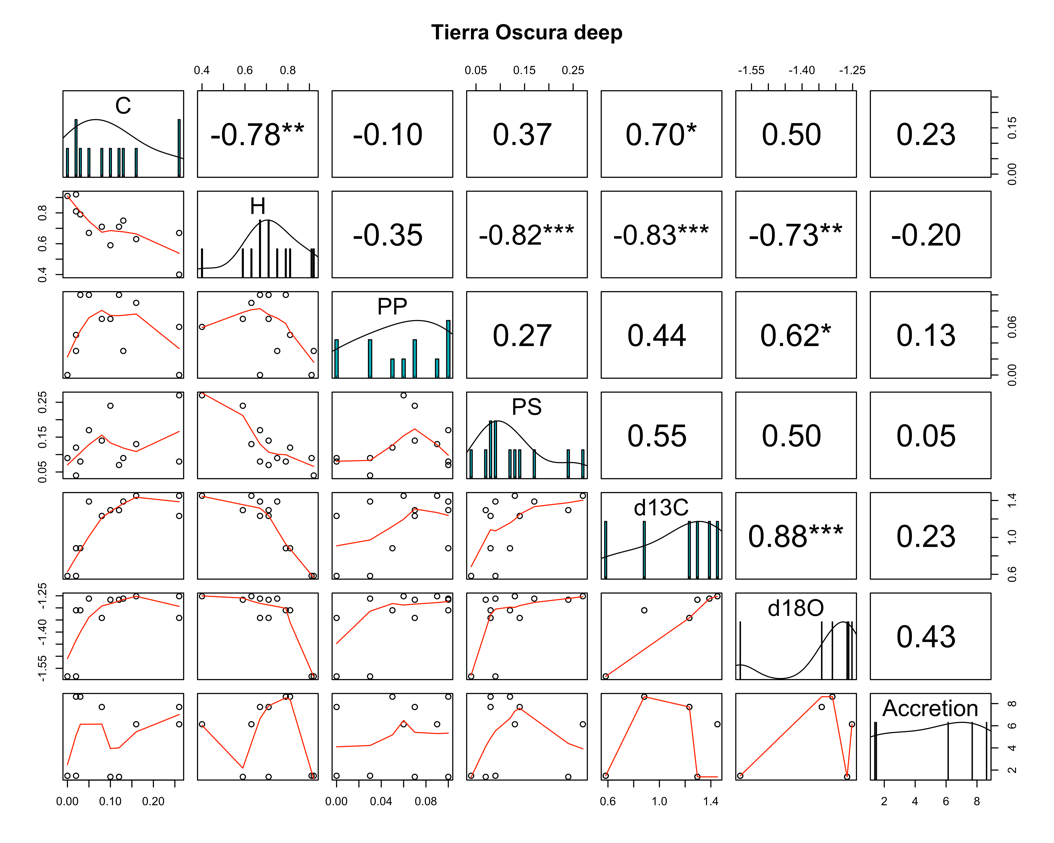

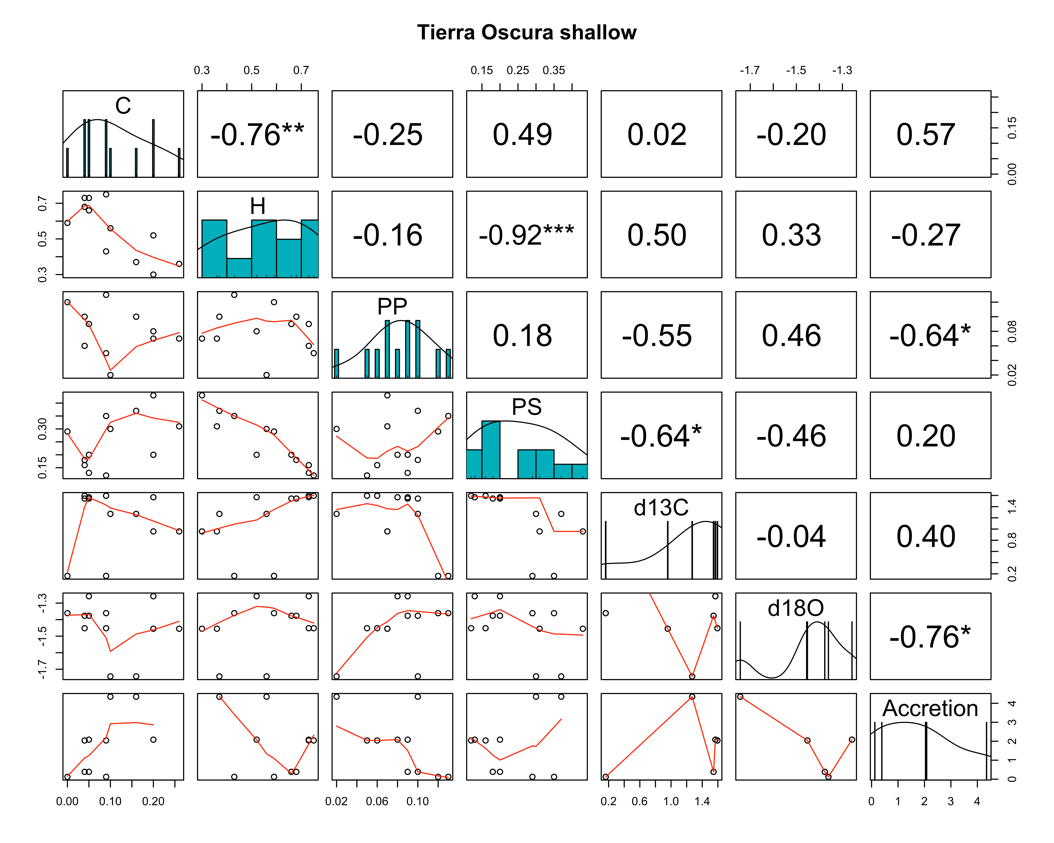

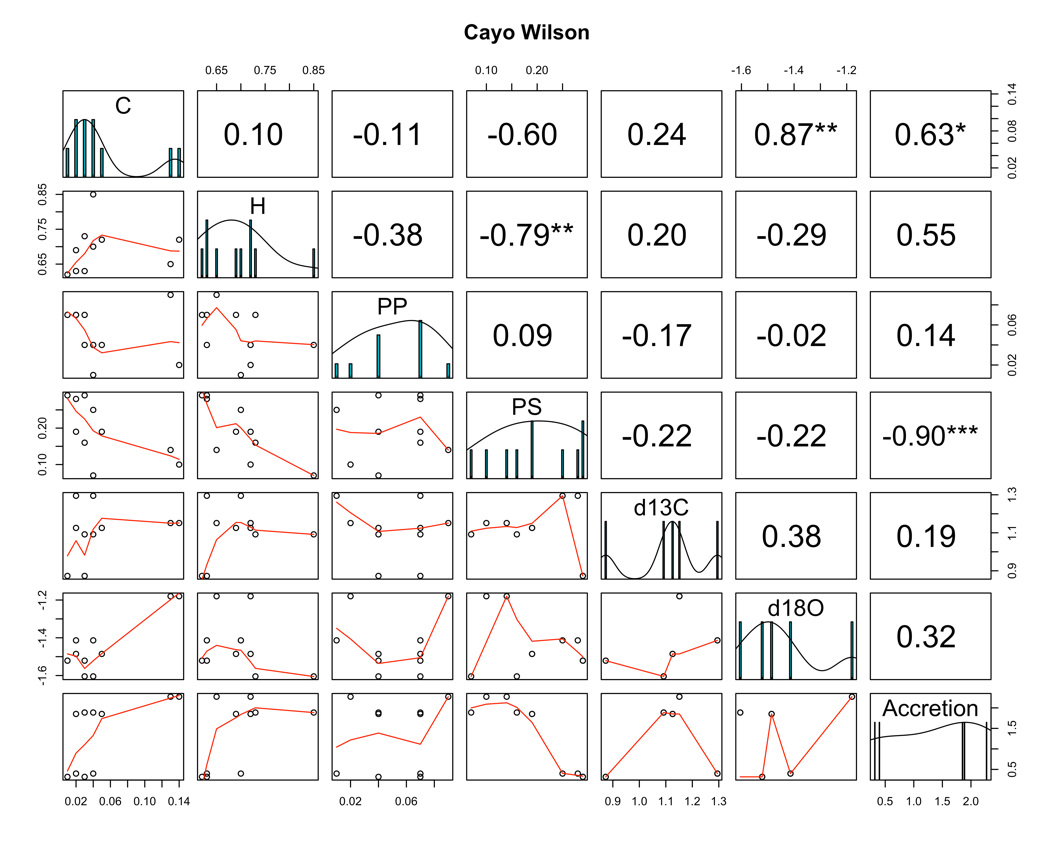

#### References

Cheng, H. et al. 2000. The half-lives of uranium-234 and thorium-230. - Chemical Geology 169: 17–33.

Clark, T. R. et al. 2014. Testing the precision and accuracy of the U–Th chronometer for dating coral mortality events in the last 100 years. - Quaternary Geochronology 23: 35–45.

Cramer, K. L. et al. 2015. Molluscan subfossil assemblages reveal the long-term deterioration of coral reef environments in Caribbean Panama. - Marine Pollution Bulletin 96: 176–187.

Cramer, K. L. et al. 2017. Prehistorical and historical declines in Caribbean coral reef accretion rates driven by loss of parrotfish. - Nat Commun 8: 14160.

Cramer, K. L. et al. 2020. Millennial‐scale change in the structure of a Caribbean reef ecosystem and the role of human and natural disturbance. - Ecography 43: 283–293.

Fredston-Hermann, A. L. et al. 2013. Marked ecological shifts in seagrass and reef molluscan communities since the mid-Holocene in the southwestern Caribbean. - BMS 89: 983–1002.

Grossman, E. L. 1987. Stable isotopes in modern benthic foraminifera; a study of vital effect. - Journal of Foraminiferal Research 17: 48–61.

Kidwell, S. M. 2013. Time-averaging and fidelity of modern death assemblages: building a taphonomic foundation for conservation palaeobiology. - Palaeontology 56: 487–522.

Kitting, C. 1985. Adaptive significance of shore-range diel migration differences in a seagrassmeadow snail population. - Contributions in Marine Science 27: 227—43.

Ludwig, K. R. 2012. Isoplot/Ex Version 3.75, a Geochronological Toolkit for Microsoft Excel. - Berkeley Geochronology Center.

Todd, J. A. 2001. Introduction to molluscan life habits database. London, UK: the Natural History Museum. Available from: https://nmita.rsmas.miami.edu/database/mollusc/mollusclifestyles.htm.

Tunnell, J. W. 2010. Encyclopedia of Texas seashells: identification, ecology, distribution, and history. - Texas A&M University Press.
